# Comparative Transcriptomic Analysis of Embryonic Stem Cells (ESCs) across Mammalian Species

**DOI:** 10.1101/2025.03.11.642582

**Authors:** Yifei Fang, Yue Su, Richard Meisel, James R Walters, Tony Gamble, Eric C. Randolph, Xingtan Yu, Meihong Shi, Guangsheng Li, Jingzhi Zhang, IISAGE Consortium, Xiuchun (Cindy) Tian, Yong Tang, Jingyue (Ellie) Duan

**Author notes:** IISAGE Consortium: Nicole C Riddle, Peggy R Biga, Anne M Bronikowski, Gerald S Wilkinson, Erica Larschan, Ritambhara Singh, Ashley Webb. Corresponding author: Xiuchun (Cindy) Tian Yong Tang Jingyue (Ellie) Duan. **Jingyue (Ellie) Duan**. **Authors’ email addresses:**. Xiuchun (Cindy) Tian. Jingyue (Ellie) Duan.

## Abstract

Pluripotency, the ability of cells to self-renew and differentiate into all the cell types in an animal’s body, is vital for mammalian early development. This study presented a comprehensive comparative transcriptomic analysis of embryonic stem cells across multiple mammalian species, defining their progression through expanded/extended, naïve, formative, and primed pluripotency states. Our findings revealed both conserved and species-specific mechanisms underlying pluripotency regulation. We also emphasized the limitations of existing state-specific markers and their limited cross-species applicability, while identifying *de novo* pluripotency markers that can inform future research. Despite variability in gene expression dynamics, gene co-expression networks showed remarkable conservation across species. Among pluripotency states, the primed state demonstrated the highest conservation, evidenced by shared markers, preserved gene networks, and stronger selective pressures acting on its genes. These findings provide critical insights into the evolution and regulation of pluripotency, laying a foundation for refining stem cell models to enhance their translational potential in regenerative medicine, agriculture, and conservation biology.

## Introduction

Embryonic stem cells (ESCs) are unique in their ability to self-renew and plasticity to differentiate into diverse cell types, offering a powerful *in vitro* model for studying developmental biology and advancing regenerative medicine^1^. These ESCs exhibit pluripotent states with distinct differentiation potentials and molecular profiles corresponding to cells of the specific stages of early embryonic development^2^. For example, mouse ESCs were initially established in the naïve pluripotency state (nESCs), representing the inner cell mass of the pre- implantation stage embryos^3^. Subsequent research has derived additional pluripotency states, including the primed state (epiblast stem cells, EpiSCs), which corresponds to post-implantation stage cells^4^, and the formative state (formative stem cells, FSCs), defined as the intermediate transition between naïve and primed states^5^. More recently, the expanded or extended state (extended pluripotency stem cells, EPSCs) has been established, representing an earlier embryonic stage with broader developmental lineage potential, such as the ability to generate both embryonic and extraembryonic tissues, thereby positioning them as pluripotent cells with certain totipotent-like characteristics^6^. Each pluripotency state is characterized by distinct differentiation potential and functional genomic profiles, serving as *in vitro* models to map ESC properties to specific embryonic stages and uncover the molecular mechanisms underlying the early embryonic programming.

Although these pluripotency states are well-characterized in the mouse model, generating ESCs at equivalent states in other mammals, such as domestic animals like pigs, cattle, and sheep, remains challenging. Specifically, the naïve state exhibits significant cross-species variability in culture conditions^7^. In contrast, the maintenance of the primed state is more consistent across species. For instance, WNT inhibition has been shown to effectively sustain EpiSCs in a range of species, including humans^8^, rats^9^, cattle^10^, pigs^11^, sheep^11^, and crab-eating macaques^12^. Moreover, comparative studies have identified conserved signaling pathways that stabilize the maintenance of a primed pluripotency state across species, such as elevating FGF and TGF pathways while inhibiting WNT signaling^13^. In contrast, the maintenance of naïve pluripotency relies on species-specific signaling mechanisms. For instance, TGF-β signaling is dispensable for maintaining mouse naïve ESCs, but it is necessary for sustaining naïve pluripotency in human cells^14^. The activation of distinct pathways highlights fundamental differences in the molecular regulation of pluripotency across species, which likely arise as evolutionary divergence associated to early developmental programming^15^. These differences are shaped by species-specific requirements for embryogenesis, such as differences in implantation strategies, developmental rates, and embryo-maternal interactions^16^. Such species-specific differences complicate the application of ESC derivation protocols across species and highlight the need to develop methodologies for identifying both universal and species-specific molecular regulatory mechanisms underlying the developmental and functional features of ESCs.

Historically, ESC lines have often been derived independently by different research groups, each focusing on a single species or a specific pluripotency state. This has limited opportunities for direct cross-species and cross-pluripotency state comparisons. Recent advances in comparative transcriptomics, supported by the availability of RNA sequencing (RNA-seq) data from various species and pluripotency states, now enable systematic investigation into conserved and species-specific gene expression patterns and regulatory mechanisms in ESCs. Comparative transcriptomics can facilitate the identification of core regulatory networks, explore evolutionary differences in gene expression, and discover new state-specific pluripotency markers that may function universally across species or exhibit distinct species-specific roles in stem cell biology.

In this study, we conducted a comprehensive comparative transcriptomics analysis of four ESC pluripotency states across nine mammalian species (three primates, four ungulates, and two rodents), using transcriptomic data from 28 research projects. We identified both common and species-specific differentially expressed genes (DEGs), enriched GO pathways between pluripotent states, assessed known state-specific markers, and discovered *de novo* pluripotency markers for each species. To further explore conserved and divergent gene regulation, we employed weighted gene co-expression network analysis (WGCNA) to construct species-specific gene networks and performed preservation analysis to identify networks conserved across species within specific pluripotency states. Additionally, our transcriptome-based evolutionary analysis revealed varying levels of selective constraints across different pluripotency states. Overall, these findings provide critical insights into the evolutionary conservation and divergence of gene regulatory mechanisms underlying pluripotency, offering a foundation for future studies to refine species-specific stem cell models and enhance cross-species applications in regenerative medicine and developmental biology.

## Results

### Transcriptome integration and online RNA-seq datasets normalization

To achieve a comprehensive analysis, we curated 120 publicly available RNA-seq samples from 28 research projects, including four ESC types, EPSCs, nESCs, FSCs, and EpiSCs, across nine mammalian species: human, mouse, rat, marmoset, crab-eating macaque, cattle, pig, sheep, and horse (**Figures 1A and 1B, Table S1**). The pluripotency states of each sample were characterized in its original research based on criteria such as ESC colony morphology, self-renewal capacity, differentiation potential, *in vitro/in vivo* developmental capability, and surface marker expression.

**Figure 1.**
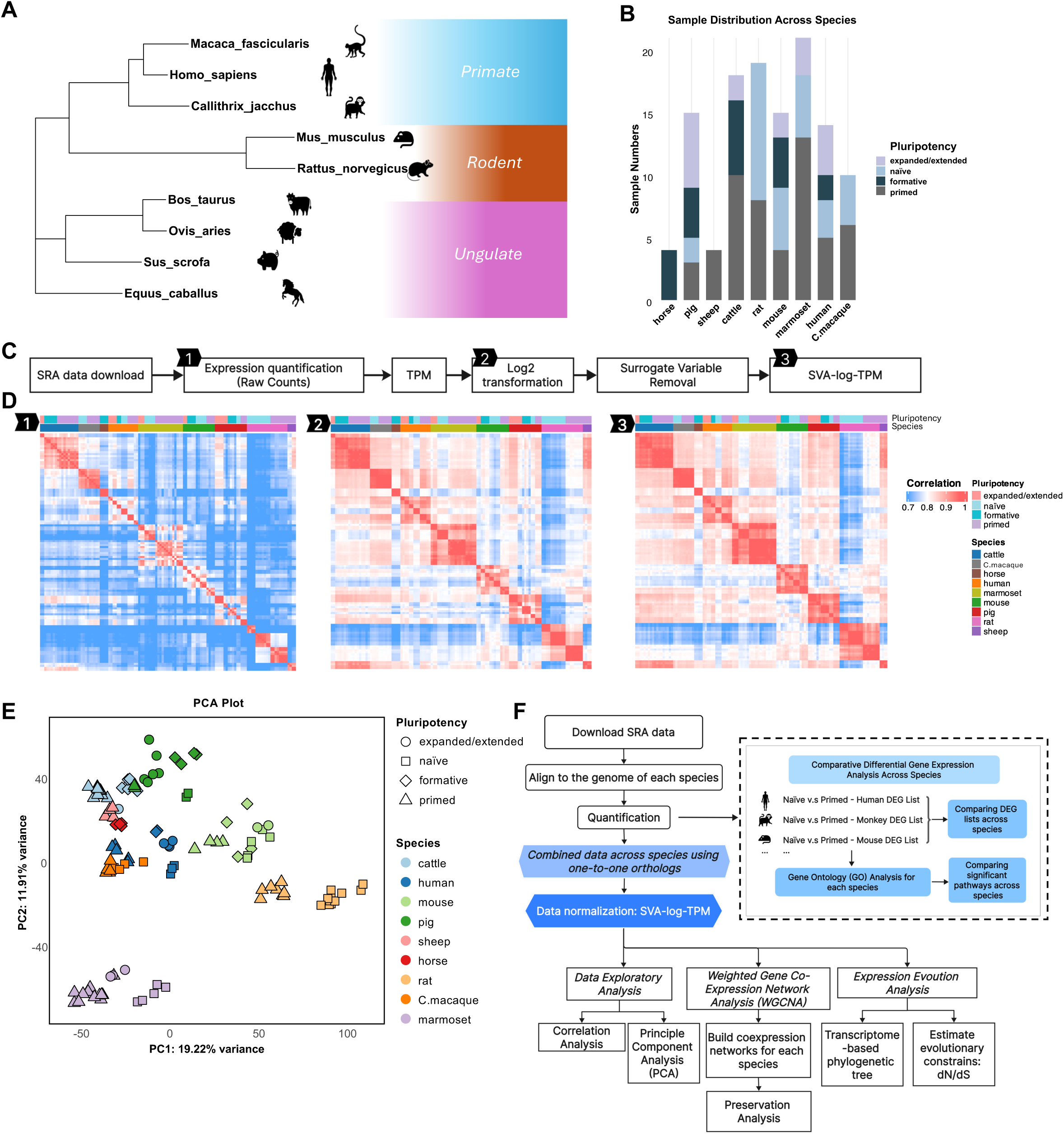
Overview of dataset integration and analytical pipeline. **A.** Phylogenetic tree representing the nine mammalian species included in the analysis. **B.** Distribution of embryonic stem cell samples with varying pluripotency levels across species. **C.** Data normalization workflow: Raw RNA-seq data were downloaded from the GEO database and processed through a unified pipeline, including alignment and quantification to obtain raw counts. TPM values were then calculated and log-transformed. To account for hidden batch effects and technical biases, Surrogate Variable Analysis (SVA) was applied, generating SVA-log-TPM values for downstream analysis. **D.** Pearson correlation coefficients comparing samples at different stages of normalization: 1) raw counts, 2) log2-TPM values, and 3) SVA-log-TPM values post-surrogate variable adjustment. **E.** Principal Component Analysis (PCA) of all samples based on SVA-log-TPM values; colors denote species, and shapes indicate pluripotency states. **F.** Detailed bioinformatics pipeline used in this study.

All raw transcriptome data were processed using a uniform bioinformatics pipeline (**Figure 1C, Table S1**). Specifically, RNA-seq reads were first aligned to species-specific reference genomes (**Table S2**), and expression read counts were quantified. To enable cross-species comparative analysis, these samples were subsequently combined using the 9,942 1-to-1 orthologous genes (i.e. genes that have only one ortholog copy among studied species) across the nine mammals. These RNA-seq data originate from diverse research projects and may be influenced by differences in sample conditions, library construction, sequencing procedures, and other unknown factors. To detect and correct for such factors, including unrecognized "hidden" biases, we applied log_2_-transformed transcripts per million (log_2_ TPM) normalization, followed by surrogate variable analysis (SVA-log-TPM) (**Figure 1C**). This SVA-log-TPM normalization greatly improved the Pearson correlation within samples of the same pluripotency state in each species, regardless of their research project origins (**Figure 1D**), confirming the reliability of our bioinformatic pipeline in handling online curated datasets for analyzing ESC transcriptomic patterns.

Principal Component Analysis (PCA) revealed that RNA-seq samples primarily clustered by species rather than pluripotency states (**Figure 1E**), reflecting a stronger species effect than the pluripotency states. These findings align with previous *in vivo* studies showing that cross-species embryo samples from pre- and post-implantation stages cluster more strongly by species than by developmental stages^17^. Variance partitioning analysis confirmed that species divergence contributes more to transcriptional variances in PCA than pluripotency states among these samples (**Figure S1A**). We further identified highly variable pluripotency-specific genes, such as *DUSP6* and *SPRY1* (**Figure S1B**), which are key components of the ERK^18^ and FGF^19^ signaling pathways and are known to play critical roles in maintaining stem cell pluripotency. Notably,

*DUSP6* has also been recognized as a novel pluripotency-related gene associated with the transition from naïve to primed states through comparative studies of mammalian embryos^17^. Moreover, our PCA results showed a consistent pattern of naïve-to-primed distribution along PC1 was observed within each species cluster (**Figure 1E**), highlighting the conservation of transcription dynamics across pluripotency stages among mammalian species.

To determine the impact of ESC derivation protocols on ESC transcriptome, we compared porcine and murine ESCs with their corresponding *in vivo* pre- or post-implantation stage embryos (**Figures S1C and S1D).** Mouse ESCs aligned well with their corresponding developmental stages: EPSC and nESC clustered closer to pre-implantation embryos, EpiSCs aligned with post-implantation embryos, and FSC, representing an intermediate state, located between pre- and post-implantation stages (**Figure S1C**). Intriguingly, porcine EpiSCs and FSCs exhibited an inverse alignment with their expected stages: EpiSCs aligned with pre-implantation embryos, while FSCs clustered closer to the post-implantation stage embryos (**Figure S1D**). We further traced back to the original studies and found that this discrepancy arose because porcine EpiSCs were derived from pre-implantation embryos^20^, whereas FSCs were derived from post-implantation embryos^21^. These results suggested that ESC transcriptomes are strongly influenced by their derivation protocols, aligning more closely with the developmental stage of their embryonic origin. Moreover, the observed impacts of derivation protocols, coupled with species-specific differences, raise limitations of applying mouse model-based practices and standards for establishing new stem cell lines across diverse species, highlighting the need to establish both universal and species-specific derivation standards and pluripotency markers.

### Gene expression comparison among pluripotency states

To minimize the impact of the inconsistencies in pluripotency characterization and assignment across species, we first identified species-specific differentially expressed genes (DEGs) during ESC pluripotency state transition by performing pairwise DEG analysis within species across four ESC states: naïve vs. primed, expanded/extended vs. naïve, naïve vs. formative, and formative vs. primed. We then compared identified DEGs and Gene Ontology (GO) biological process terms across species that had both ESC types presented (**Figure 1F and Table S3-S4**).

### nESCs vs. EpiSCs comparison

We compared DEGs between the nESCs and EpiSCs across six species: rat, marmoset, macaque, pig, mouse, and humans (**Figure 2**). Our results showed that the species-specific DEGs dominated the upregulated DEGs in nESCs, with 1,637 in mice, 1,474 in rats, and 1,343 in humans (**Figure 2A)**. Despite this species specificity, we identified two DEGs, *ZBTB10* and *IHO1,* that were consistently upregulated in nESCs across all six species (**Figure 2A**). *ZBTB10*, a transcription factor (TF), interacts with core pluripotency factors *OCT4* and *SOX2,* playing a role in maintaining stem cell pluripotency^22^. *IHO1* is known to promote the DNA double-strand break formation^23^, a crucial mechanism for maintaining ESC genome stability^24^.

**Figure 2.**
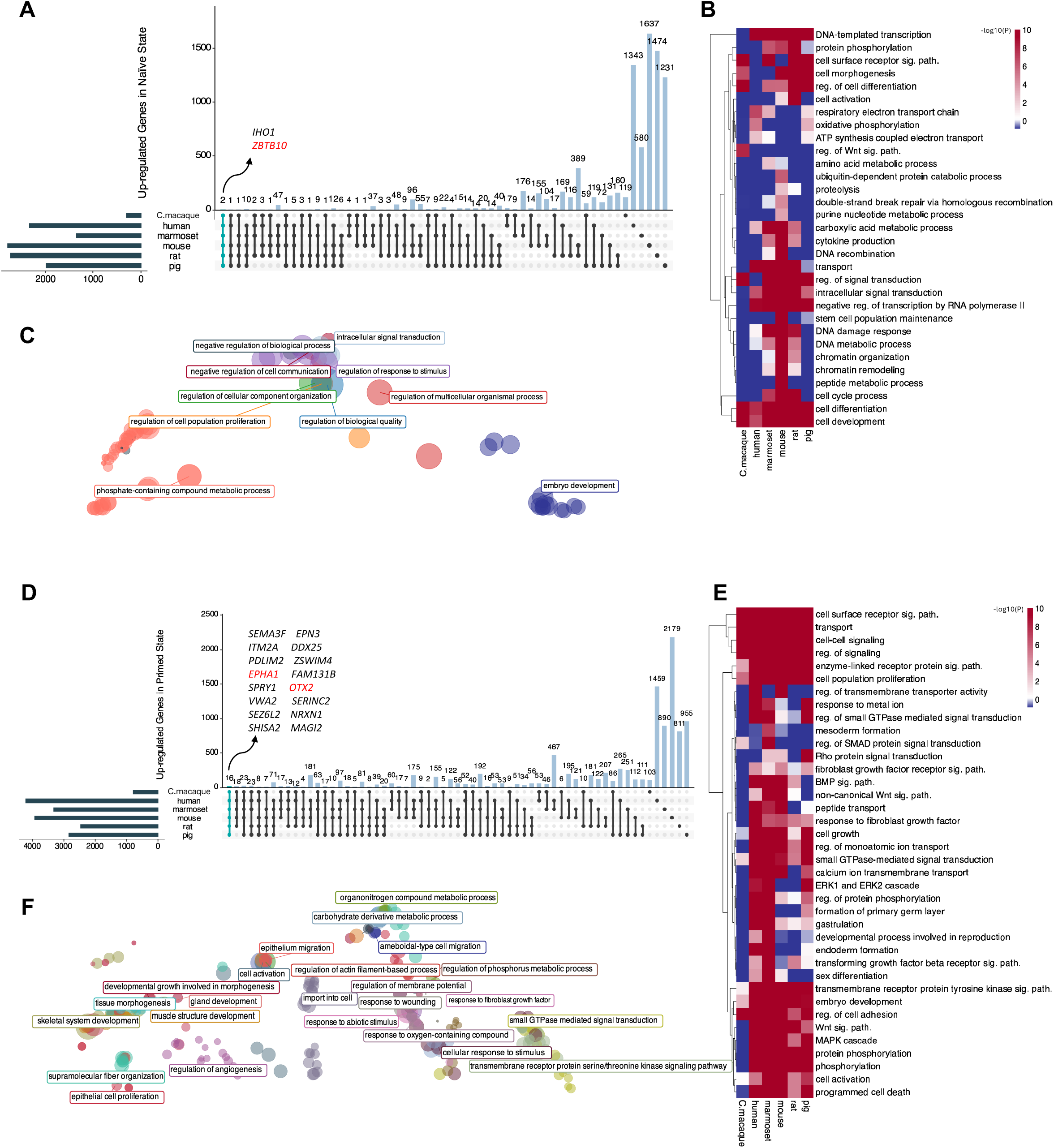
Comparative analysis of shared and species-specific genes and GO pathways in naïve versus primed ESC states. **A.** Up-regulated genes identified in the naïve state across multiple species. **B.** Selected GO pathways significantly enriched by the up-regulated genes in the naïve state across species. **C.** All conserved GO:BP pathways significantly enriched by up-regulated genes in the naïve state across species, with terms clustered based on similarity. **D.** Up-regulated genes identified in the primed state across multiple species. **E.** Selected GO pathways significantly enriched by the up-regulated genes in the primed state across species. **F.** All conserved GO:BP pathways significantly enriched by up-regulated genes in the primed state across species, with terms clustered based on similarity.

We also observed that species with a closer evolutionary relationship shared more common DEGs. For instance, 176 nESCs upregulated DEGs were shared between humans and marmosets, while 389 DEGs overlapped between mice and rats (**Figure 2A**). GO analysis revealed significant enrichment of pathways across all species, including cell differentiation and cell development, suggesting that these pathways are fundamental to the core processes that maintain naïve pluripotency and are evolutionary conserved (**Figures 2B and 2C; Table S4**). Moreover, we identified several mouse-specific enriched pathways, including the purine nucleotide metabolic process, peptide metabolic process, ubiquitin-dependent protein catabolic process, and stem cell population maintenance (**Figure 2B**).

Similarly, species-specific DEGs also dominated the upregulated DEGs in EpiSCs compared to nESCs, with 2,179 identified in mice, 1,459 in humans, and 955 in pigs (**Figure 2D**). 16 commonly upregulated genes in EpiSCs were identified across six species, including *OTX2, EPHA1,* etc. (**Figure 2D**). *OTX2* is known to stabilize the EpiSC state by preventing neural differentiation^25^. *EPHA1*, a member of the Eph receptor tyrosine kinase family, promotes pluripotency maintenance in human ESCs and iPSCs^26^. GO analysis of EpiSCs upregulated DEGs revealed significantly enriched pathways for primed state ESCs across species, such as non-canonical Wnt signaling, BMP signaling, ERK and MAPK cascade, transmembrane receptor protein tyrosine kinase signaling, embryo development, and regulation of cell adhesion (**Figures 2E and 2F; Table S4**). These pathways are essential for lineage specification, cellular interactions, signaling, and developmental processes in primed state ESC. Their conservation across species suggests their evolutionary importance in regulating early development and ensuring proper establishment of a robust primed pluripotency state.

### nESCs vs. EPSCs comparison

In the comparison of nESCs and EPSCs across human, marmoset, mouse, and pig (**Figure S2**), two commonly upregulated genes in nESCs, *DDIT4* and *BCL3,* were identified (**Figure S2A**). *DDIT4* mediates HIF1α and mTOR signaling, which are critical for pluripotency^27^, while *BCL3* links LIF-STAT3 to core pluripotency genes *Oct4* and *Nanog* to promote the maintenance of naïve pluripotency^28^. GO analysis revealed conserved pathways in nESCs, such as cellular developmental process, regulation of cellular and biological processes, cell differentiation, and intracellular signaling cassette (**Figures S2B and S2C**). In EPSCs, 18 commonly upregulated genes were identified, including well-known pluripotency regulators *MYC* and *DNMT3B* (**Figure S2D**). *MYC* facilitates cell reprogramming and pluripotent state establishment^29^, while *DNMT3B* supports LIF-dependent ESC self-renewal^30^. Shared GO pathways in EPSCs included cell surface receptor signaling, regulation of cell adhesion, and embryonic morphogenesis, revealing core regulatory processes of expanded/extended pluripotency state maintenance (**Figures S2E and S2F; Table S4**).

### FSCs vs. nESCs comparison

In the comparison between FSCs and nESCs across humans, mice and pigs (**Figure S3**), 140 commonly upregulated DEGs in formative states were identified. Several key genes involved in transitioning from naïve to primed pluripotency, ESC proliferation, and maintaining stem cell states including *LIN28A*^31^, *ETV4*^32^, *DUSP6*^17^, *ID3*^33^, *USP44*^34^, *MYC*^35^, and *DNMT3B*^36^ (**Figure S3A**). Enriched GO pathways in FSCs among three species included cell-cell signaling by WNT, cell differentiation, and MAPK cascade (**Figures S3B and S3C**). Conversely, 30 common upregulated genes were identified in the naïve state, such as *YPEL2, WHAMM, WDR62,* and *ULK1* (**Figure S3D**). *ULK1* is essential for mouse embryonic stem cell self-renewal and pluripotency^37^. Shared GO pathways in nESCs included regulation of cell communication, signal transduction, cell cycle, and apoptotic process (**Figures S3E and 3F; Table S4**).

### EpiSCs vs. FSCs comparison

When comparing EpiSCs to FSCs across humans, mice, pigs, and cattle (**Figure S4**), five commonly shared upregulated genes in the primed state were *MYL9, PNKD, ATP1B2*, *TSPAN7*, and *NRARP* (**Figure S4A)**. *TSPAN7* regulates pluripotency in neural stem cells and neural differentiation^38^, while *NRARP,* a part of the Notch signaling pathway, plays a key role in differentiation and pluripotency maintenance^39,40^. Shared GO terms in all species except mice included regulation of cell differentiation, cell surface receptor, and transmembrane receptor protein tyrosine kinase signaling pathways (**Figures S4B and S4C; Table S4**). In FSCs, two shared upregulated genes were *ZBTB11*, and *GDF3* (**Figure S4D**). *ZBTB11* maintains pluripotency by repressing pro-differentiation genes in ESCs^22^. *GDF3*, a TGF-beta superfamily member, regulates ESCs pluripotency through BMP inhibition in a species-specific manner^41^. Enriched pathways in FSCs across four species involved regulation of gene expression, and DNA-templated transcription (**Figure S4E and S4F**).

### Evaluation of known and novel pluripotency state-specific markers

Next, we evaluated the performance of known pluripotency state-specific markers (**Table S5**) defined in mice to assess their expression levels and applicability across mammals (**Figure 3**). Our analysis revealed that while these state-specific markers effectively differentiated pluripotency states in mouse ESCs, they failed to consistently distinguish their corresponding pluripotency states across all species (**Figure 3A-D)**. For example, expanded/extended markers (n=8) did not show the highest expression of EPSCs compared to other ESC states in humans (**Figure 3A**). Similarly, naïve markers (n=7) generally performed well across species except for differentiating nESC from EpiSCs in macaque (**Figure 3B**). Moreover, formative (n=14) and primed (n=9) markers were able to differentiate FSCs and EpiSCs from EPSCs or nESCs but could not distinguish between each other in most species that contained these two states (**Figures 3C and 3D**).

**Figure 3.**
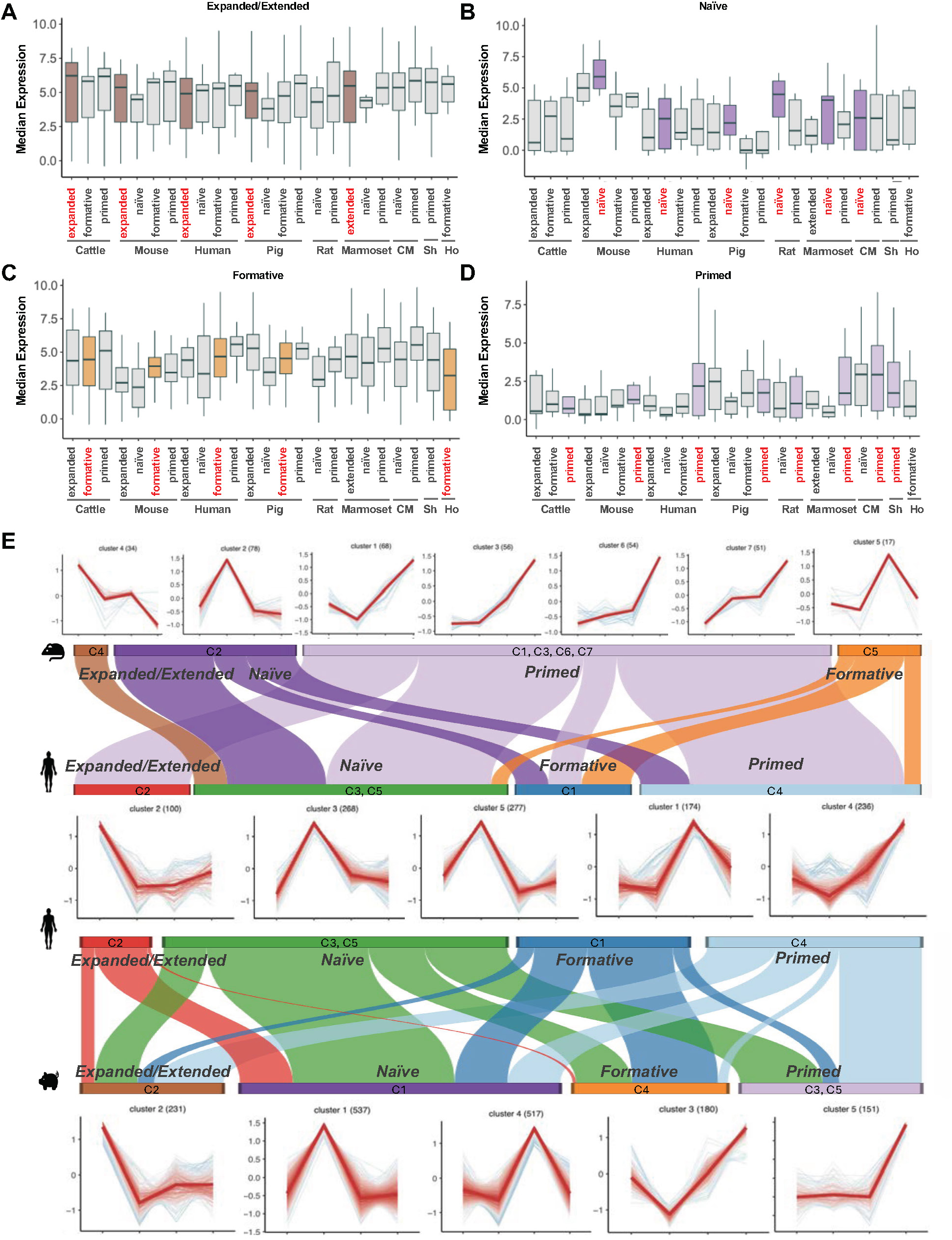
Assessment of known state-specific markers across species and identification of de-novo markers. **A-D**. Evaluation of known mouse state-specific markers for the expanded/extended state (A), naïve state (B), formative state (C), and primed state (D). CM represents crab-eating macaque. Sh represents sheep and Ho represents horse. **E.** *de novo* state-specific markers identified across species. Markers were determined through differential expression analysis and subsequently clustered using the mfuzz method to trace their shared expression trajectory changes across the four pluripotency states. The comparison of shared and species-specific markers between mouse and human is displayed at the top, while the comparison between human and pig is shown at the bottom.

Given the limitations of mouse-based ESC markers, we aim to identify *de novo* state-specific markers applicable across multiple mammalian species. Using pairwise DEGs from species with all four pluripotency states represented (humans, mice, and pigs), we identified genes uniquely upregulated in one specific state per species and assigned them to the corresponding state cluster (**Figure 3E**). We applied soft clustering (see Methods) to identify the common expression trajectories of these markers across states within each species (**Figure S5A-B**). Comparing the clusters between human, mouse, and pig orthologs, we identified genes assigned to the same or different clusters of pluripotency states (**Figure 3E**).

Common genes assigned in the same state-specific clusters across species were further defined as state-specific markers. For example, *CHRNA4,* a potential regulator of pluripotency maintenance^42^, was identified as the only primed state-specific marker shared across all three species (**Figure 4A**). Other state-specific markers were significantly upregulated in their respective states in at least two species, such as *DBNDD1* in expanded/extended state, *BHMT*^43^ in naïve state, and *SNORD17*^44^ in formative state. Some of these genes play known roles in pluripotency-related functions and show consistently elevated expression in their corresponding states across species (**Figure 4A**). In contrast, pluripotency-specific genes that did not overlap across species were identified as species- and state-specific genes (**Figures 4B, S5, and Table S6)**. Interestingly, genes such as *LEF1, SMC6,* and *LSR* were uniquely elevated in the expanded/extended state in humans, mice, or pigs, respectively, but not in the other two species, suggesting that each species possesses a unique set of markers tailored to their pluripotency regulation (**Figure 4B and Table S6)**.

**Figure 4.**
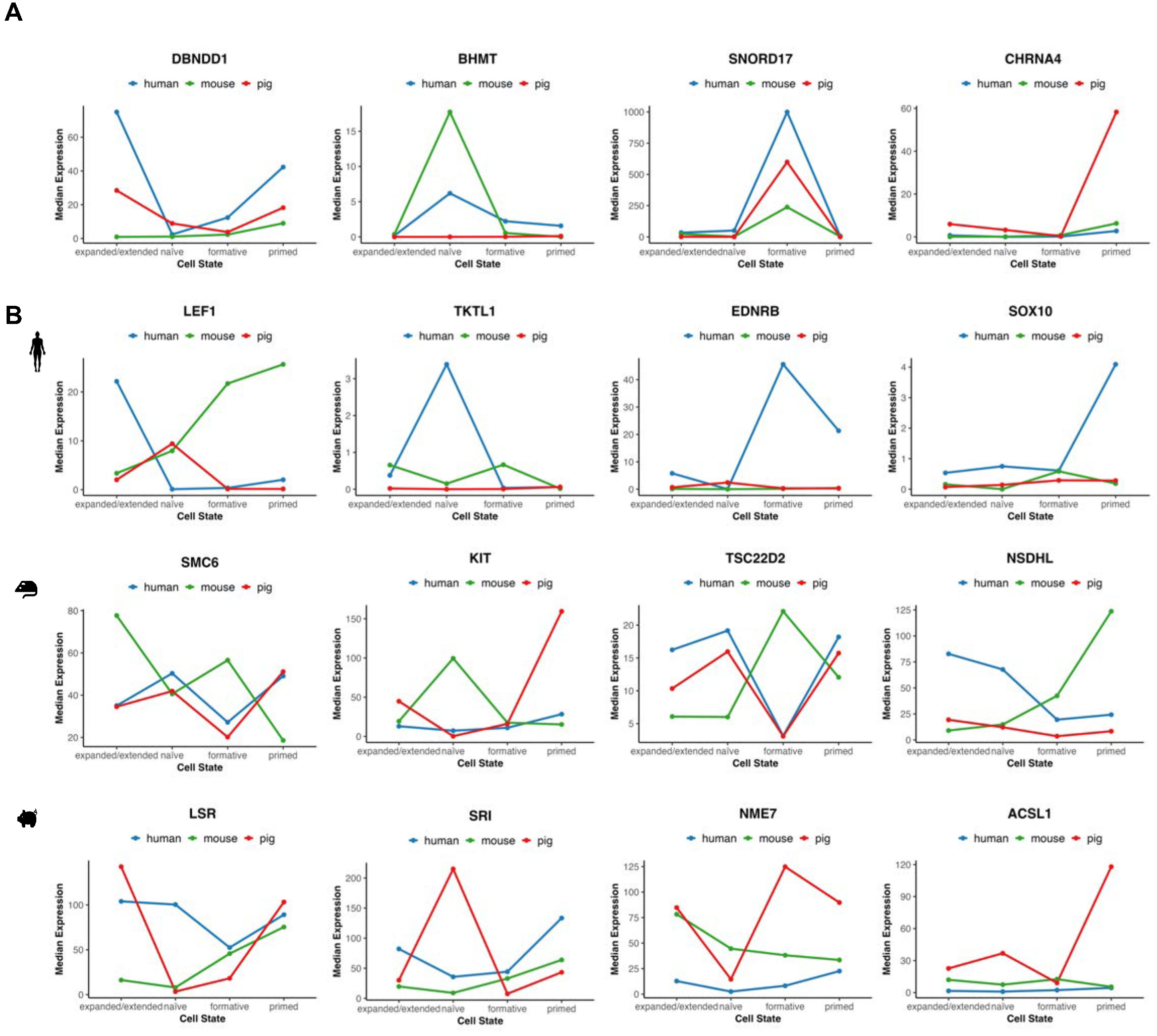
Representative examples of shared and species-specific de-novo markers. **A.** Expression trajectory changes across the four pluripotency states for shared markers between different species (e.g., *DBNDD1* for the expanded/extended state, *BHMT* for the naïve state, *SNORD17* for the formative state, and *CHRNA4* for the primed state). **B.** Expression trajectory changes across the four pluripotency states (expanded/extended; native; formative and primed) for species-specific markers. Human-specific examples are shown at the top, mouse-specific examples in the middle, and pig-specific examples at the bottom. TPM values were used.

### Gene Co-Expression Network Construction and Preservation Analysis

To further identify pluripotency state-specific gene networks and assess their conservation across species, we performed weighted gene co-expression network analysis (WGCNA) across the four pluripotency states in all species except sheep, horses, and macaque due to insufficient sample sizes (**Figures 5 and S6**). This analysis identified several modules significantly correlated with each specific pluripotency state. For example, in the human pluripotency network, the modules labeled salmon (R = 0.87), turquoise (R = 0.96), tan (R = 0.95), black (R = 0.94), and blue (R = 0.85) were significantly associated with the expanded/extended, naïve, formative and primed states, respectively (**Figure 5A**). Similarly, in mice, the modules labeled lightgreen (R= 0.85) and blue (R= 0.8) were significantly correlated with the primed state in mice **(Figure 5B)**, while in pigs, modules labeled green (R= 0.89) and pink (R= 0.91) were enriched in primed state (**Figure S7**).

**Figure 5.**
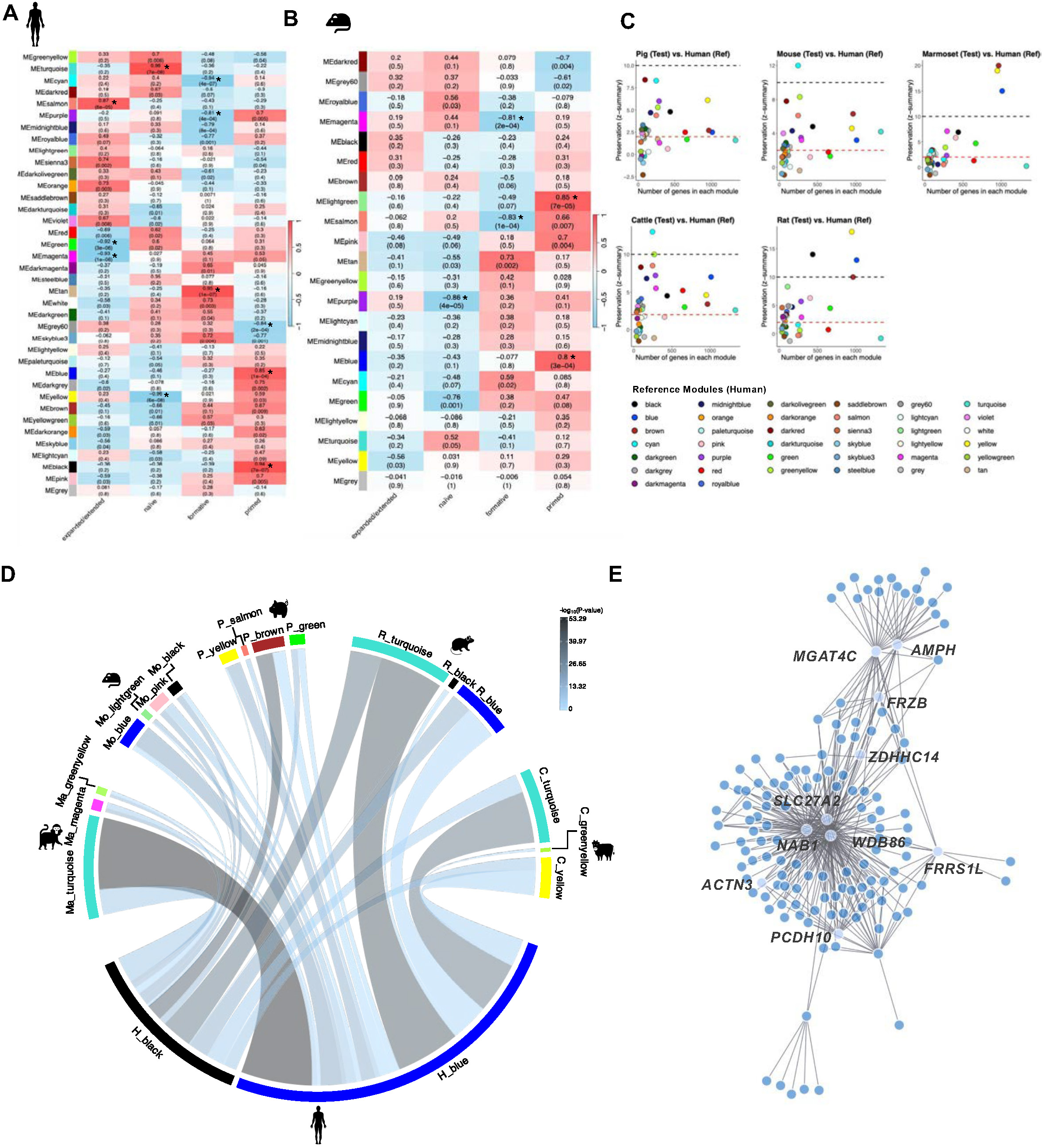
Cross-species comparison of WGCNA analysis. **A.** Module-trait relationships within the human co-expression network, with modules showing a correlation coefficient >= 0.8 and p < 0.05 marked by an asterisk. **B.** Module-trait relationships within the mouse co-expression network, highlighting modules with a correlation coefficient > 0.8 and p < 0.05 by an asterisk. **C.** Z-statistics depicting module preservation of the human network (reference network) in networks of other species. Thresholds for weak preservation (Z = 2) and strong preservation (Z = 10) are indicated in the plots. **D.** The top two modules from other species’ networks where the human blue and black modules are mainly preserved. The thickness of the connecting bands represents the number of overlapping genes between the two modules, while the color indicates the –log (P) value, where P represents the statistical significance of the overlap. **E.** Co-expression network of the human blue module, with the top 10 hub genes (ranked by centrality scores using Kleinberg’s metric) labeled.

To assess the conservation of pluripotency state-specific gene modules across species, we conducted a preservation analysis using the human network as a reference (**Figures 5C, 5D and Figure S8**). This analysis identified several conserved modules between humans and another species, where a module with a Z-summary score > 2 indicated weak preservation and a Z-summary score > 10 reflects strong preservation. Notably, the blue, black, yellow, brown, purple, and darkred modules from the human network were preserved across all species analyzed (**Figure 5C**).

Given that the blue and black modules had a strong correlation with the human primed state, we further investigated whether the preserved modules in other species also correlated with this state. We found that the human blue module was largely preserved within the blue, light-green, and pink modules in mice (**Figure 5D**), and turquoise modules of marmosets, cattle, and rats. All of these modules showed significant correlations to the primed state in their respective species (**Figures S7B-D**). However, in pigs, the human blue module was preserved in the yellow and salmon modules (**Figure 5C**), which were significantly enriched in the extended/expanded state (**Figure S7A)**. Similarly, the human black module, also associated with the primed state, was substantially preserved in modules such as pink in mice, brown and green in pigs, turquoise and blue in rats, and turquoise and yellow in cattle **(Figure 5D** and **Figure S7)**. These preserved modules exhibited significant primed state associations in their respective species, except for the magenta and greenyellow modules in marmoset and black modules in mice, which lacked strong primed correlations (**Figure 5D** and **Figure S7C**). These findings indicate that the human blue and black modules represented conserved gene co-expression networks associated with the primed state across most mammalian species examined, with some species-specific exceptions.

We further visualized the human gene network of these conserved modules (**Figures 5E and S9A**) and identified several key hub genes: *NAB1*, a transcriptional repressor^45^, along with *SLC27A2*, *MGAT4C*, and *AMPH*, involved in lipid metabolism^46^, glycosylation^47^, and endocytosis^48^, respectively, were identified as key hub genes in the blue module. Similarly, *ELAPOR2*, associated with the regulation of autophagy and apoptosis^49^, *LTA4H*, a mediator of leukotriene-associated inflammation^50^, and *TSPAN7*, contributing to cell adhesion, migration, and neural development^51^, were identified as key hub genes in the black module. GO analysis further revealed pathways related to neuronal development, tissue morphogenesis, and system differentiation (**Figures S9B and S9C**), indicating that primed ESCs are poised for lineage commitment, consistent with their role in later embryonic development stages.

### Transcriptome-based phylogenies

To gain insights into the evolution of gene expression, we reconstructed transcriptome-based phylogenetic trees for four distinct pluripotency states (**Figure 6A**). Each state exhibited a unique tree structure, reflecting dynamic transcriptional changes and interspecies variations across different pluripotency states. Rodents and primates consistently clustered closely across all states, whereas ungulates displayed a more varied clustering pattern. For example, porcine FSCs formed a distinct branch rather than clustering with cattle and horses in the formative-specific phylogenetic tree (**Figure 6A**).

**Figure 6.**
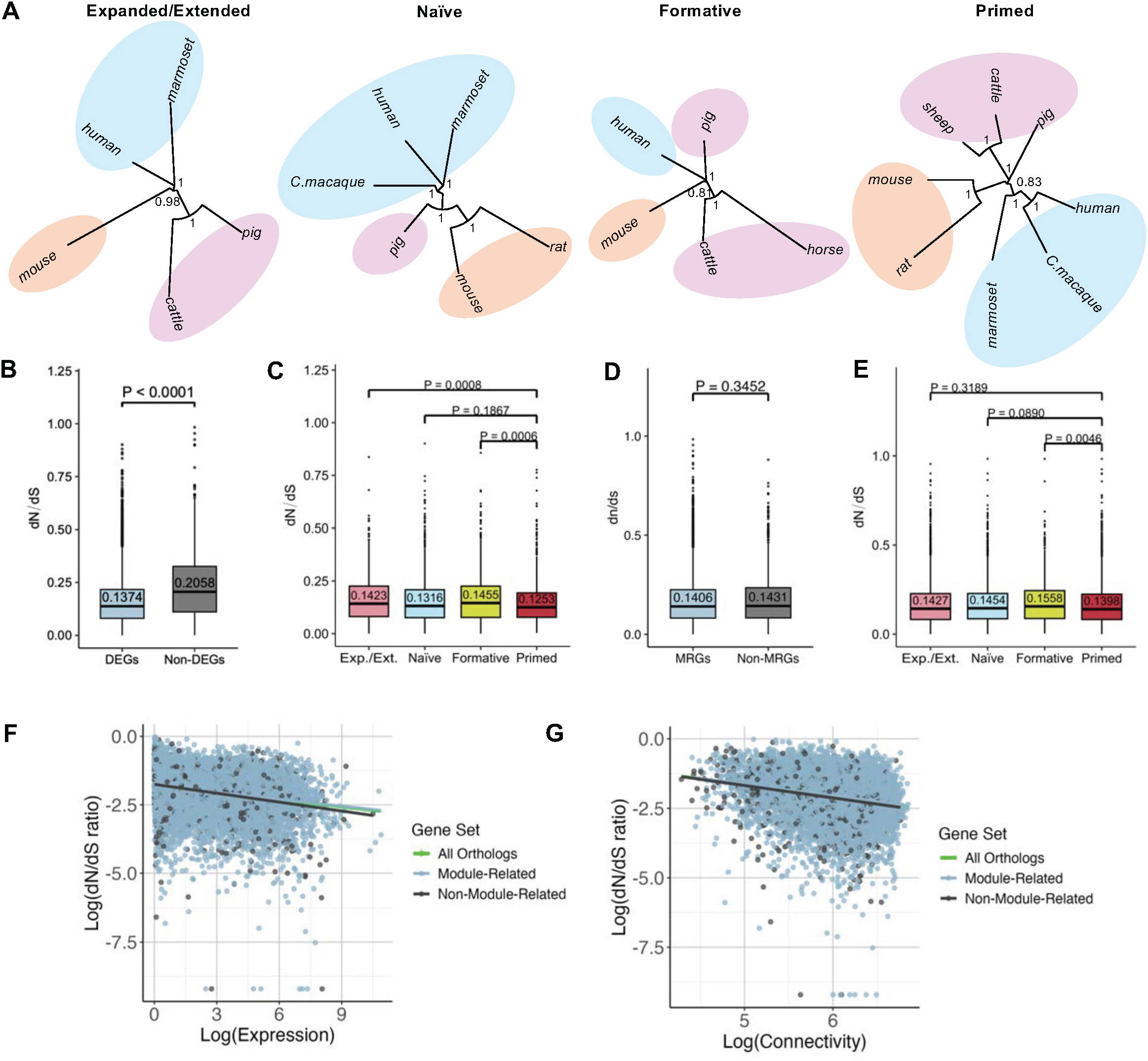
Analysis of expression evolution. **A.** Transcriptome-based phylogenetic trees for the four pluripotency states. Neighbor-joining trees were constructed using 1:1 orthologous genes, with branch lengths representing the proportion of expression variation corresponding to evolutionary expression changes. Bootstrap values (calculated from 1,000 random resamplings of 1:1 orthologous genes) are indicated. **B-E.** Box plots illustrating dN/dS ratios for various gene sets: DEGs versus non-DEGs (B), state-specific genes identified through DEG analysis (C), module-associated genes (MRGs) identified by WGCNA as related to any pluripotency state versus Non-MRGs (D), and genes in WGCNA modules associated with specific states (E). P-values above each comparison indicate the significance of dN/dS differences between gene sets (Mann–Whitney U test). **F-G.** dN/dS ratios as functions of (F) gene expression and (G) connectivity identified by WGCNA. Smoothing functions represent different gene sets, with the green line for all orthologous genes, the blue line for module-associated genes (MRGs), and the black line for non-module-associated genes (Non-MRGs). Negative correlations are observed between gene expression and dN/dS ratio, as well as between connectivity and dN/dS ratio.

Among all states, the primed state showed the closest alignment with known mammalian phylogeny based on DNA or protein sequences, suggesting it represents a conserved post-implantation development stage, whereas other states exhibit more developmental diversity or species-specific variations (**Figure 6A**). This is further supported by the consistent use of FGF2 and Activin-A (F/A) in the culture media for EpiSCs across species, in contrast to more variable supplements used for naïve conditions between mice and humans^13^.

### Evolutionary rates are correlated with pluripotency

To investigate whether evolutionary rates of protein genes are associated with pluripotency states, we calculated the dN/dS ratio, which indicates the rate of change in amino acid sequences compared to neutral changes in a protein-coding gene^52^. DEGs identified from pairwise pluripotency state comparisons exhibited a significantly lower dN/dS ratio (0.14) compared to non-DEGs (0.21, P < 0.0001, Mann–Whitney U test, **Figures 6B and S10A**). While both median dN/dS values were below 1, indicating purifying selection, DEGs had a significantly lower dN/dS ratio, suggesting stronger evolutionary constraints. However, since DEGs also showed higher expression levels than non-DEGs, we cannot fully exclude gene expression as a confounding factor in the observed evolutionary rate differences.

To ensure these results were not biased by a large number of total DEGs, we further divided the DEG and non-DEG by species (**Figure S10E**) or separated DEGs in specific pairwise comparisons (expanded extended/naïve, naïve/formative, formative/primed) within each species (**Figure S10F).** Across comparisons, DEGs consistently showed significantly lower dN/dS values than non-DEGs, except for the Mouse formative/primed comparison (**Figure S10F**). These findings reinforce that genes involving pluripotency state transitions are generally under stronger purifying selection with a slower evolutionary rate^53^.

We further analyzed dN/dS ratios for pluripotency-specific genes uniquely expressed in different pluripotency states among all species. Our results revealed that the primed state had the significantly (*P* < 0.05, Mann–Whitney U test) lowest dN/dS ratios compared to other states (**Figures 6C and S10B**), suggesting higher conservation of the primed states related genes with slower evolutionary rates. Similarly, genes from WGCNA modules (MRGs) correlated with specific states exhibited lower dN/dS ratios than those from non-correlated modules (Non-MRGs, **Figures 6D and S10C**). Among these modules, the primed state showed the lowest dN/dS values (**Figures 6E and S10D**). Finally, we observed strong negative correlations between gene expression levels and dN/dS ratios (*P* <1e-10, linear model, **Figure 6F**), as well as between WGCNA module connectivity and dN/dS ratios (*P* < 1e-10, linear model, **Figure 6G**). To further investigate the relationships among these variables, we performed a correlation and partial correlation analysis to assess how dN/dS, gene expression levels, and connectivity interact (**Table S7**). The results confirmed that genes with higher expression levels or greater connectivity in gene co-expression networks tend to evolve more slowly, consistent with the hypothesis that highly expressed and functionally central genes are subject to stronger purifying selection. These findings indicate that genes were highly expressed or had more gene network connections evolve more slowly. Overall, these results highlight the close relationship between evolutionary rates and pluripotency across states, with genes associated with pluripotency transitions, particularly in the primed state, showing stronger evolutionary constraints and slower rates of evolution.

## Discussion

Pluripotency is a fundamental biological process regulated by intricate and complex gene regulatory networks. Embryonic stem cells (ESCs) exist in distinct pluripotency states, each corresponding to specific stages of early embryonic development and characterized by unique molecular profiles. To better understand the molecular divergence underlying these states, we performed a comprehensive comparative transcriptomic analysis to investigate the gene expression dynamics during pluripotency states transition across mammalian species. Our findings reveal both conserved and species-specific mechanisms of pluripotency regulation associated with different pluripotency states while highlighting the evolutionary constraints shaping their regulation.

### Dynamics of gene expression in pluripotency states transition

Cross-species DEG analysis revealed a predominance of species-specific DEGs, with only a few conserved genes such as *ZBTB10* and *IHO1* in the naïve state and *OTX2* in the primed state. *ZBTB10*, a C2H2 zinc finger BTB domain TFs expressed in human and mouse early embryos, is downregulated upon differentiation^54^. Although *ZBTB10* knockdown has minimal effects on ESC morphology or OCT4-positive cell numbers, it may have accessory or self-renewal functions^22^.

Conversely, the functions of *IHO1* in pluripotency remain underexplored, underscoring the need for further investigation. *OTX2,* a key transcription factor facilitating the transition from naïve to primed state, has been widely recognized as essential for maintaining primed pluripotency^55,25,56^. GO enrichment analysis revealed that key signaling pathways of pluripotency, including the Wnt, BMP, and FGF, are conserved in the primed state across mammalian species. However, how these pathways further interact with species-specific regulatory networks remains an open question.

To capture intermediate dynamics of the pluripotency transition, we incorporated the formative state between naïve and primed states. This analysis identified conserved upregulated genes including *LIN28A*, *DNMT3B*, *ETV4*, and *DUSP6* during the naïve-to-formative transition, while *ZBTB11* was downregulated during the formative-to-primed transition. These findings align with previous studies. For example, *LIN28A*^5^, *DNMT3B*^57^, and *ETV4*^58^ are well-documented formative markers in mice. while *DUSP6* is known to associate with the naïve-to-primed transition^17^ and up-regulated in the intermediate state of human PSCs^59^. *ZBTB11* has also previously been identified as an essential transcription factor for maintaining pluripotency^22^. Further GO analysis highlighted consistently conserved pathways across several states, such as the MAPK cascade which has been shown to play a pivotal role in balancing proliferation and differentiation in stem cells^60^. Additionally, genes upregulated in the formative state were enriched in pathways related to DNA recombination and chromosome organization. These pathways may be linked to X-chromosome inactivation (XCI), a hallmark event during the transition between naïve and primed states, which involves extensive chromatin reorganization^13,61^.

### Identification of pluripotency state-specific markers and co-expression hub genes

Our analysis revealed that mouse pluripotency state-specific markers were insufficient to distinguish pluripotency states in other species. We further identified conserved and species-specific *de novo* markers for each pluripotent state. We observed that the primed state had the highest number of shared markers, with *CHRNA4* conserved across humans, mice, and pigs. *CHRNA4* encodes the neuronal acetylcholine receptor subunit alpha-4 (nAChRα4) and is involved in neural differentiation^62^. Our findings revealed substantial variability in species-specific gene expression patterns and uncovered an intriguing phenomenon that a state-specific cluster in one species aligned with a different state cluster in another species. This indicates the need for caution when selecting or applying markers from one species to another in cross-species analyses.

While state-specific genes are poorly conserved across species, the structure of WGCNA co-expression networks appear to be well-preserved among mammals, consistent with previous comparative transcriptomic analyses involving data from eight tissues, such as skin, mammary gland, and marrow^63^. This conservation may reflect the integrated nature of pluripotency regulation, which relies on interactions between various signaling pathways rather than isolated gene activity^64^. In our analysis, several human co-expression primed-specific modules were consistently preserved in cattle, pigs, mice, rats, and marmosets. Notably, the human blue module and its preserved counterparts in other species showed a significant correlation with the primed state across species. Key hub genes in this module include *NAB1*, a transcription coregulator involved in nervous system development^45^, and *SLC27A2*, which is essential for lipid biosynthesis and fatty acid metabolism by activating long-chain fatty acids into their CoA derivatives^46^. Further GO analysis highlighted enrichment of pathways associated with neuronal development and differentiation, supporting the notion that cells in the primed state are poised for lineage commitment while retaining elements of the pluripotency network^65^.

### Primed state exhibits the highest evolutionary conservation

We further calculated the dN/dS ratios for DEGs, non-DEGs, and state-specific upregulated genes. Our results revealed that DEGs generally have lower dN/dS values than non-DEGs, suggesting that genes involved in pluripotency regulation are subject to stronger purifying selection. Such evolutionary constraints align with our observation that the core pluripotency regulatory network demonstrates significant conservation, indicating the critical role of pluripotency-related genes in maintaining essential cellular processes^66^. Interestingly, primed state-specific genes exhibited significantly lower dN/dS values compared to genes specific to other states, indicating intensified selective pressure on this state. This aligns with the primed state showing the highest evolutionary conservation, as demonstrated by transcriptome-based phylogenetic clustering, a greater number of shared DEGs, and a higher WGCNA module preservation across species. This conservation may be due to the consistent derivation of EpiSCs from the embryonic epiblast cells of the post-implantation embryonic stage across species, in contrast to nESCs, which are derived from the more variable pre-implantation stage. Moreover, the uniformity of culture conditions for EpiSCs across species may contribute to the transcriptomic similarities, in contrast to the species-specific conditions required for naïve states^13^. Additionally, a previous study^67^ comparing several pre- and post-implantation stage embryos of monkeys, pigs, and humans revealed that post-implantation samples clustered closer across species than pre-implantation samples in PCA. This finding suggests that the observed conservation of the primed state primarily reflects the intrinsic evolutionary stability of post-implantation epiblast cells during early development, supporting the idea of the hourglass model, which proposes that embryonic divergence is highest at the earliest and latest stages, while the mid-embryonic (phylotypic) stage exhibits the greatest evolutionary conservation^68^.

## Conclusion

Our study provides a comparative transcriptomic analysis across mammalian species, revealing both conserved and species-specific mechanisms of pluripotency regulation. The primed state emerged as the most evolutionarily conserved state, with shared pathways and gene networks highlighting its critical role in preparing cells for lineage commitment. While species-specific gene expression patterns highlight evolutionary diversity, the preservation of gene co-expression network structures suggests a broader regulatory network conserved across species. The knowledge gained here provides a foundation for standardizing pluripotency state characterization within species and optimizing ESC culture systems across diverse species. These advancements hold the potential to enhance applications in regenerative medicine, sustainable agriculture, and conservation while deepening our understanding of the evolutionary mechanisms shaping early mammalian development.

## METHOD DETAILS

### Curation and quantification of transcriptome data

A total of 120 RNA-seq samples from embryonic stem cells (ESCs) at various pluripotent and totipotent states (expanded or extended, naïve, formative, and primed) were collected across nine species: human, mouse, rat, marmoset, crab-eating macaque, cattle, pig, sheep, and horse. These samples, derived from 28 studies, were sourced from the NCBI GEO database (https://www.ncbi.nlm.nih.gov/geo/). The specific references for the RNA-seq data are provided, and detailed information for all samples is summarized in Supplementary Table S1. For certain species, RNA-seq data for specific pluripotency stages were unavailable in public databases (last searched on January 1, 2024), and these cell types were excluded from further analysis.

All samples underwent a standardized upstream analysis pipeline. The RNA-seq data were downloaded using the SRA Toolkit v3.0.5 (https://github.com/ncbi/sra-tools/wiki/01.-Downloading-SRA-Toolkit) with the fastq-dump function. Quality control was performed using fastp v0.23.4^69^ with the parameters: -f 15 -F 15 -q 20 -u 40 -n 8 -l 30. Subsequently, alignment was carried out with STAR v2.7.10b,^70^ mapping each sample to its corresponding genome (Supplementary Table S2) using the parameters: --quantMode TranscriptomeSAM -- outSAMtype BAM SortedByCoordinate --outSAMunmapped Within --readFilesCommand zcat. Finally, RSEM v1.3.3^71^ was used for gene-level quantification with default settings.

### Ortholog retrieval and data normalization

We retrieved the one-to-one orthologs for the species in our dataset from the Ensembl database release 109^72^, identifying 9,941 orthologous genes with 1:1 orthology relationships among all the species studied. The RNA-seq samples from different species were integrated using these orthologous genes, enabling cross-species comparisons based on the merged data. TPM normalization was performed, with values calculated using RSEM, and subsequently transformed to log2 (N + 1) values (log2-TPM). To detect and correct for hidden biases like those arising from variations in sampling conditions and sequencing protocols across different experiments, we employed the SVA R package (v3.50.0)^73^. This approach generated SVA-log2-TPM values, which were subsequently used for further analysis.

### Data exploratory analysis

Principal Component Analysis (PCA) was performed using the prcomp function in R, based on the SVA-log2-TPM values. Pearson correlation analysis was conducted on the raw counts, log2-TPM values, and SVA-log2-TPM values, both before and after normalization. Additionally, variance partitioning analysis was applied to quantify the contributions of species and pluripotency state to gene expression variance for each gene. This analysis was implemented using the variancePartition R package (v1.32.3)^74^, with species and state as fixed effects in a linear mixed model.

### Differential gene expression analysis

We employed the DESeq2 R package (v1.42.0)^75^ to identify differentially expressed genes (DEGs) in each species by comparing nESCs to EpiSCs and across successive states: EPSCs to nESCs, nESCs to FSCs, and FSCs to EpiSCs. To control for multiple hypothesis testing and improve statistical power, we applied Independent Hypothesis Weighting^76^. Genes exhibiting an absolute log2 fold-change greater than 1 and an adjusted p-value less than 0.05 were classified as DEGs. We then performed an overlap analysis of DEGs across different species to identify those that are commonly shared using the UpSetR R package (v1.4.0)^77^.

### Identification of de novo state-specific markers

State-specific markers were defined as genes significantly upregulated in a specific state compared to all other states. Marker identification was first performed within each species to capture intra-species specificity. To examine expression dynamics across states, we applied Mfuzz clustering using the clusterGVis R package (v0.1.1) (https://github.com/junjunlab/ClusterGVis) to group genes into predominant expression trajectories. Subsequently, cross-species comparisons were conducted to identify both conserved markers shared across species and those unique to individual species, providing insights into universal and species-specific pluripotency regulation.

### Gene co-expression network construction and preservation analysis

We utilized the WGCNA R package (v1.72-5)^78^ to construct gene co-expression networks for each species individually. The blockwiseModules function from WGCNA in R was employed for the analysis, using SVA-log-TPM data as input. The soft-thresholding power was determined with the pickSoftThreshold function to optimize the scale-free topology fit of the network. A signed network was constructed to retain the sign of the correlation in the adjacency matrix. To evaluate whether modules were preserved across species, we conducted a preservation test using the modulePreservation function, with the human network as a reference. Z-scores were obtained to measure the preservation, with Z>10 suggesting strong preservation, 2<Z<10 representing weak preservation and Z<2 meaning no preservation^79^. Hub genes within the networks were identified based on the hub centrality scores via Kleinberg’s metric using the hub_score function from the igraph v2.0.3 package^80^. The relationships among genes within the modules were visualized using networkD3 R package (v0.4) (https://github.com/christophergandrud/networkD3).

### Gene expression phylogenies

Aligned with Brownian-motion-based models which assumes that gene-expression evolution occurs through a succession of independent changes in gene-expression levels, we employed a variance-based method to estimate expression divergence between each pair of species^81^. Distance matrices were calculated for each pluripotency state based on the SVA-log2-TPM values of all orthologous genes. Using these distance matrices, we constructed phylogenetic trees using the neighbor-joining (NJ) approach. The reliability of the branching patterns was assessed through bootstrap analyses, wherein 1:1 orthologous genes were randomly sampled with replacement 1000 times. The bootstrap values represent the proportions of replicate trees that support the branching pattern of the consensus tree displayed in the figures.

### dN/dS ratio calculation

The coding sequences (CDS) of all orthologous genes were retrieved from the Ensembl database. Multiple sequence alignments were performed using TranslatorX^82^ with MAFFT as the alignment method. For phylogenetic reconstruction, RAxML v8.2.12^83^ was used to infer maximum-likelihood gene trees. The ratio of nonsynonymous substitutions per nonsynonymous site (dN) to synonymous substitutions per synonymous site (dS) was calculated using PAML (codeml) v4.10.6^84^. A homogeneous model was applied (model = 0, NSsites = 0) to estimate the overall dN/dS ratio for each orthologous gene. Additionally, we examined the relationships between dN/dS ratios and two gene properties: gene expression levels and gene connectivity (i.e., the number of direct interactions a gene has with other genes). Gene connectivity was derived from WGCNA. To further investigate the interplay among these factors, we performed a correlation and partial correlation analysis to assess how dN/dS, gene expression, and connectivity interact. Specifically, we computed Spearman correlations for all pairwise comparisons and further estimated partial correlations to disentangle direct and indirect relationships between these factors. To ensure robustness, we applied a bootstrapping approach (1,000 replicates) to calculate confidence intervals for the partial correlations.

### Gene ontology enrichment analyses

Gene ontology analysis is conducted using gprofiler2 R package (v 0.2.3)^85^. GO lists were grouped by similar terms based on their semantic similarity to reduce redundancy by rrvgo R package (v1.14.2)^86^.

## Resource availability

### Lead contact

Requests for further information and resources should be directed to and will be fulfilled by the lead contact, Jingyue (Ellie) Duan.

### Materials availability

This study did not generate new unique reagents.

### Data and code availability

All datasets and source code used in this study: https://github.com/coderFaye/CS-ESC

## Acknowledgments

The author thanks all members of the IISAGE Consortium for the discussion of the results. This study was supported by grants from the NSF BII (#2213824) to J. E. D and the IISAGE Consortium.

## Author Contributions

Y.F., J.E.D. X.T., and Y.T. conceptualized the research. Y.F. conducted the sample curation and data analysis. Y.S. supported dataset curation. R.M., J.W., A.G., E.R., X.T., Y.T., and IISAGE Consortium contributed to the analysis methods and interpretation of the data. X.Y., M.S., G. L., and J.Z. helped with manuscript preparation and data discussion. Writing-original draft, Y.F. and J.E.D. All co-authors participated in revising the manuscript. Funding acquisition J.E.D. and the IISAGE Consortium.

## Supplementary information

Document S1. Figures S1–S10

Document S2. Tables S1-S7

Document S3. Table S4

