## Supplementary material for "Comparative Transcriptomic Analysis of Embryonic Stem Cells (ESCs) across Mammalian Species": Figure S1A

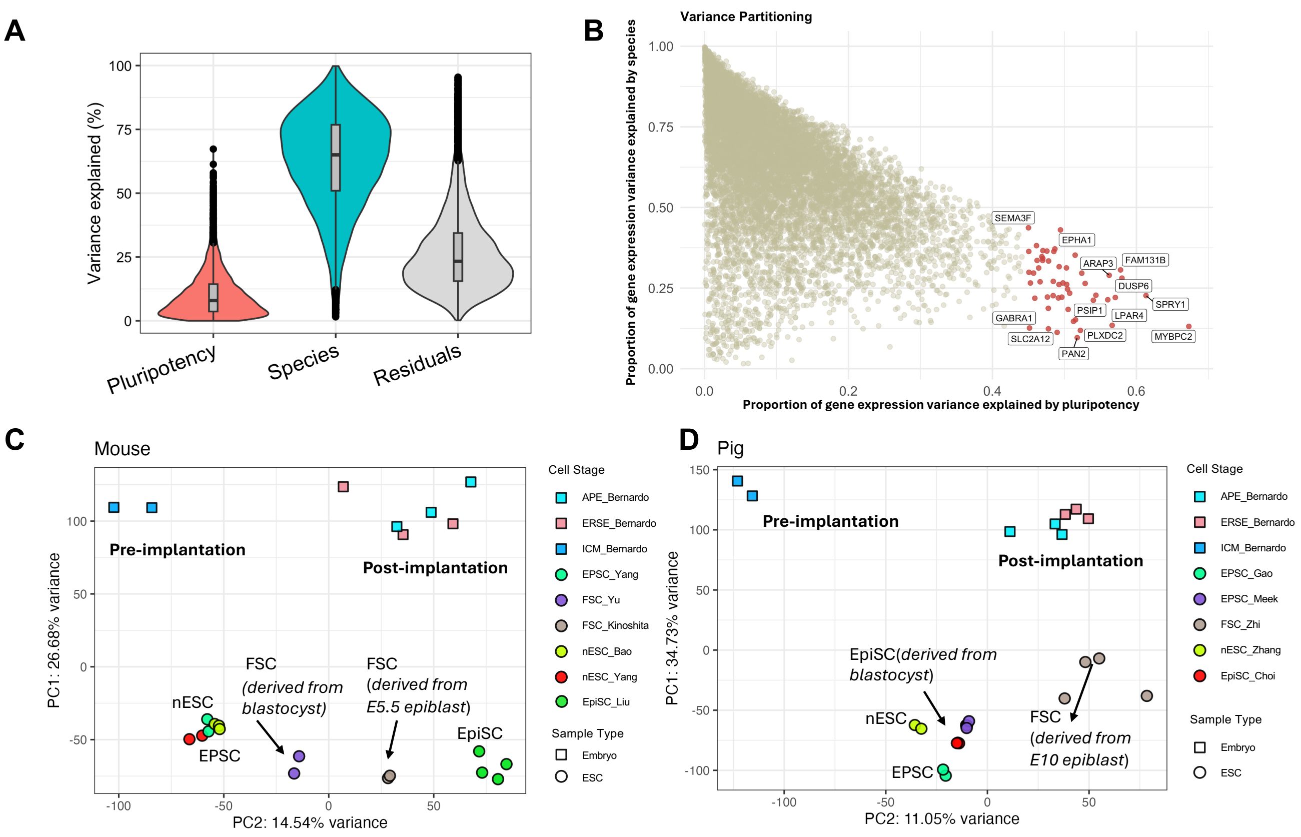


**Supplementary Figure 1. Variance partitioning, gene-specific variance, and PCA analysis linking ESC samples to in vivo embryonic stages.**

**A.** Variance partitioning analysis across all samples, displaying the contribution of two key variables: pluripotency state and species.

**B.** Gene-specific variance partitioning, with genes showing the highest variance attributed to pluripotency state highlighted.

**C.** PCA analysis mapping all the porcine ESC samples to in vivo embryo samples. Embryo samples are classified into three developmental stages: ICM (inner cell mass, pre-implantation), ERSE (epithelial radially symmetric epiblast, post-implantation), and APE (anterior-posterior epiblast, post-implantation). ESC samples from distinct studies are color-coded, and sample types (embryo or ESC) are represented by distinct shapes.

**D.** PCA plot showing the alignment of mouse ESC samples with in vivo embryo samples, categorized into the same three developmental stages: ICM, ERSE, and APE.

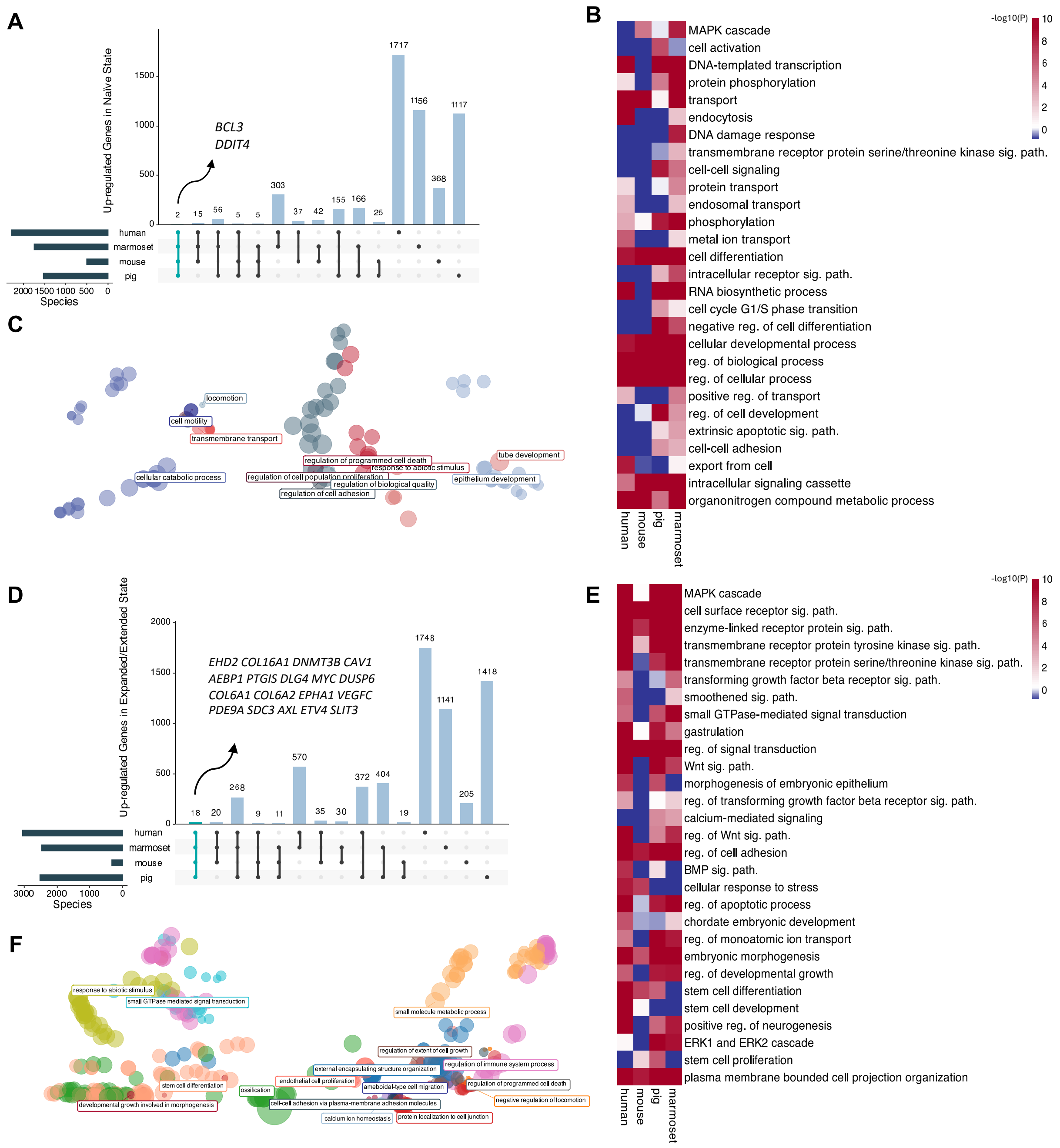


**Supplementary Figure 2. Comparative analysis of shared and species-specific genes and GO pathways in expanded/extended versus naïve ESC states.**

**A.** Up-regulated genes identified in the naïve state across multiple species.

**B.** Selected conserved and species-specific Gene Ontology (GO) pathways significantly enriched by the up-regulated genes in the naïve state across species.

**C.** All conserved GO biological process (GO:BP) pathways significantly enriched by up-regulated genes in the naïve state across species, with terms clustered based on similarity.
**D.** Up-regulated genes identified in the expanded/extended state across multiple species.
**E.** Selected conserved and species-specific GO pathways significantly enriched by the up-regulated genes in the expanded/extended state across species.
**F.** All conserved GO:BP pathways significantly enriched by up-regulated genes in the expanded/extended state across species, with terms clustered based on similarity.

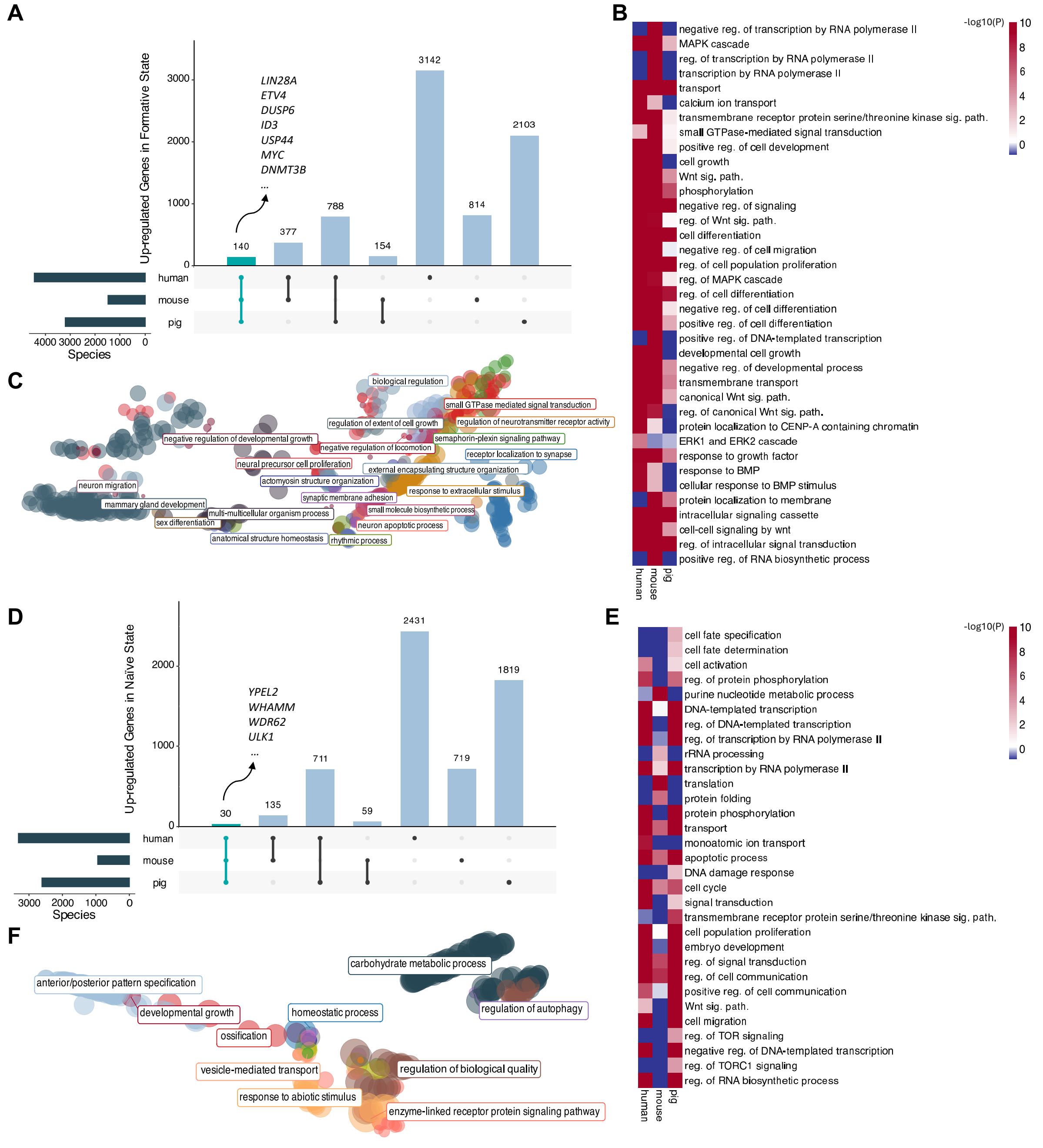


**Supplementary Figure 3. Comparative analysis of shared and species-specific genes and GO pathways in naïve versus formative ESC states.**

**A.** Up-regulated genes identified in the formative state across multiple species.

**B.** Selected conserved and species-specific Gene Ontology (GO) pathways significantly enriched by the up-regulated genes in the formative state across species.

**C.** All conserved GO:BP pathways significantly enriched by up-regulated genes in the formative state across species, with terms clustered based on similarity.

**D.** Up-regulated genes identified in the naïve state across multiple species.

**E.** Selected conserved and species-specific Gene Ontology (GO) pathways significantly enriched by the up-regulated genes in the naïve state across species.

**F**. All conserved GO:BP pathways significantly enriched by up-regulated genes in the naïve state across species, with terms clustered based on similarity.

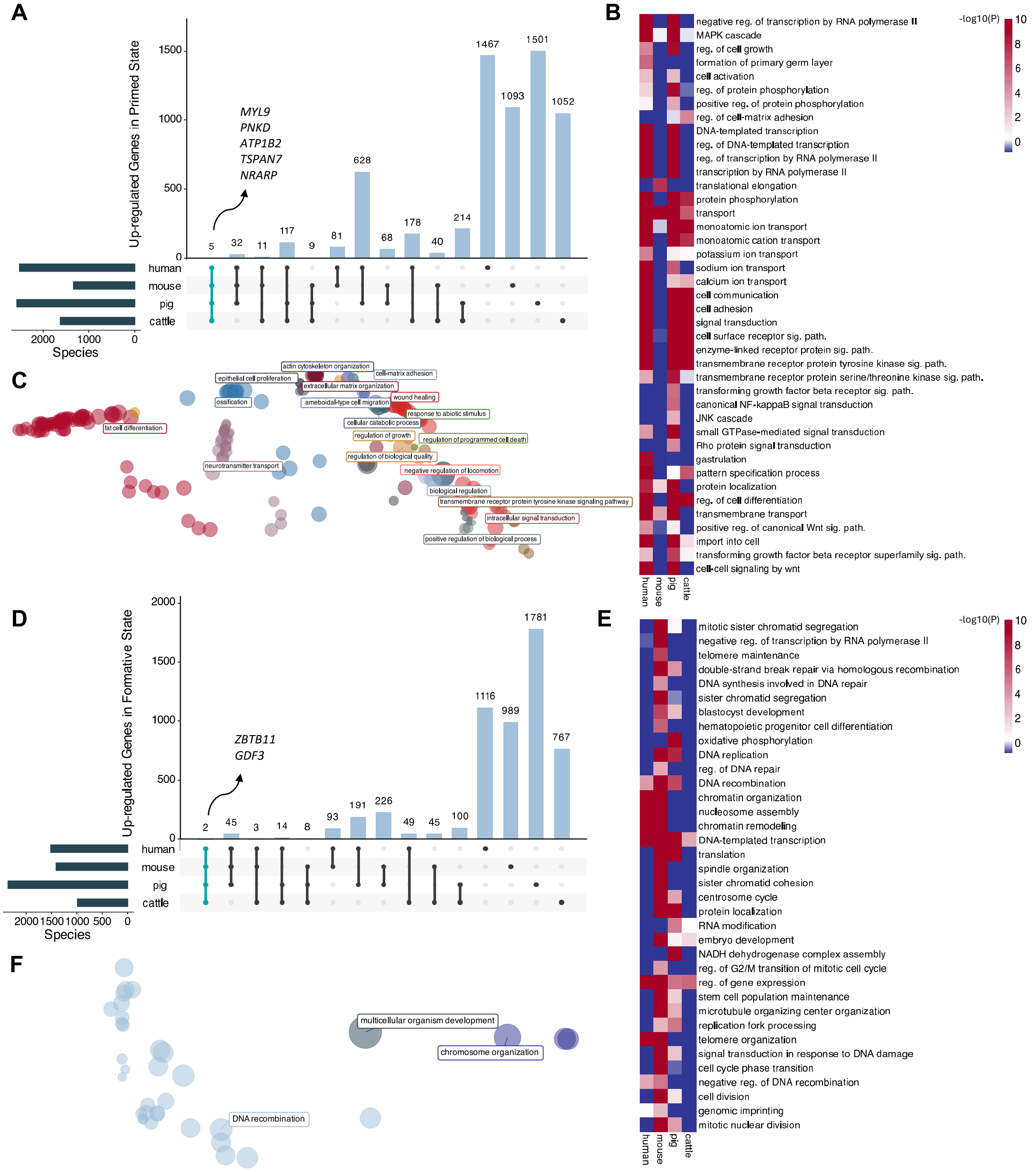


**Supplementary Figure 4. Comparative analysis of shared and species-specific genes and GO pathways in formative versus primed ESC states.**

**A.** Up-regulated genes identified in the primed state across multiple species.

**B.** Selected conserved and species-specific GO pathways significantly enriched by the up-regulated genes in the primed state across species.

**C.** All conserved GO biological process (GO:BP) pathways significantly enriched by up-

regulated genes in the primed state across species, with terms clustered based on similarity.
**D.** Up-regulated genes identified in the formative state across multiple species.
**E.** Selected conserved and species-specific GO pathways significantly enriched by the up-regulated genes in the formative state across species.
**F.** All conserved GO biological process (GO:BP) pathways significantly enriched by up-regulated genes in the formative state across species, with terms clustered based on similarity.

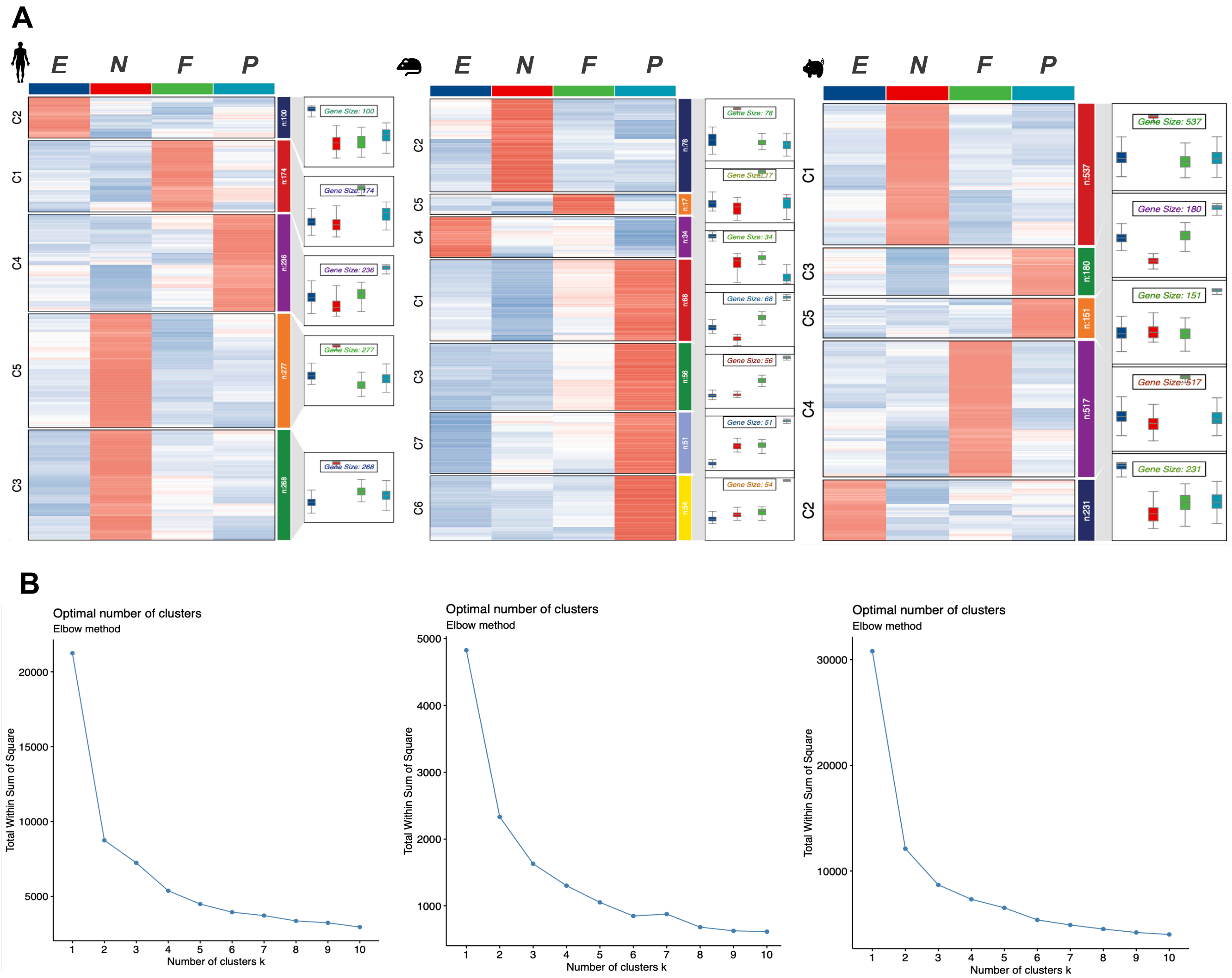


**Supplementary Figure 5. Identification and clustering of de-novo state-specific markers across human, mouse, and pig ESCs.**

**A.** Heatmaps and boxplots displaying de-novo state-specific markers for human (left), mouse (center), and pig (right).

**B.** Elbow plots indicating the optimal number of clusters for mfuzz clustering in human (left), mouse (center), and pig (right). The selected clusters for subsequent analysis were determined based on both the elbow plots and their biological relevance.

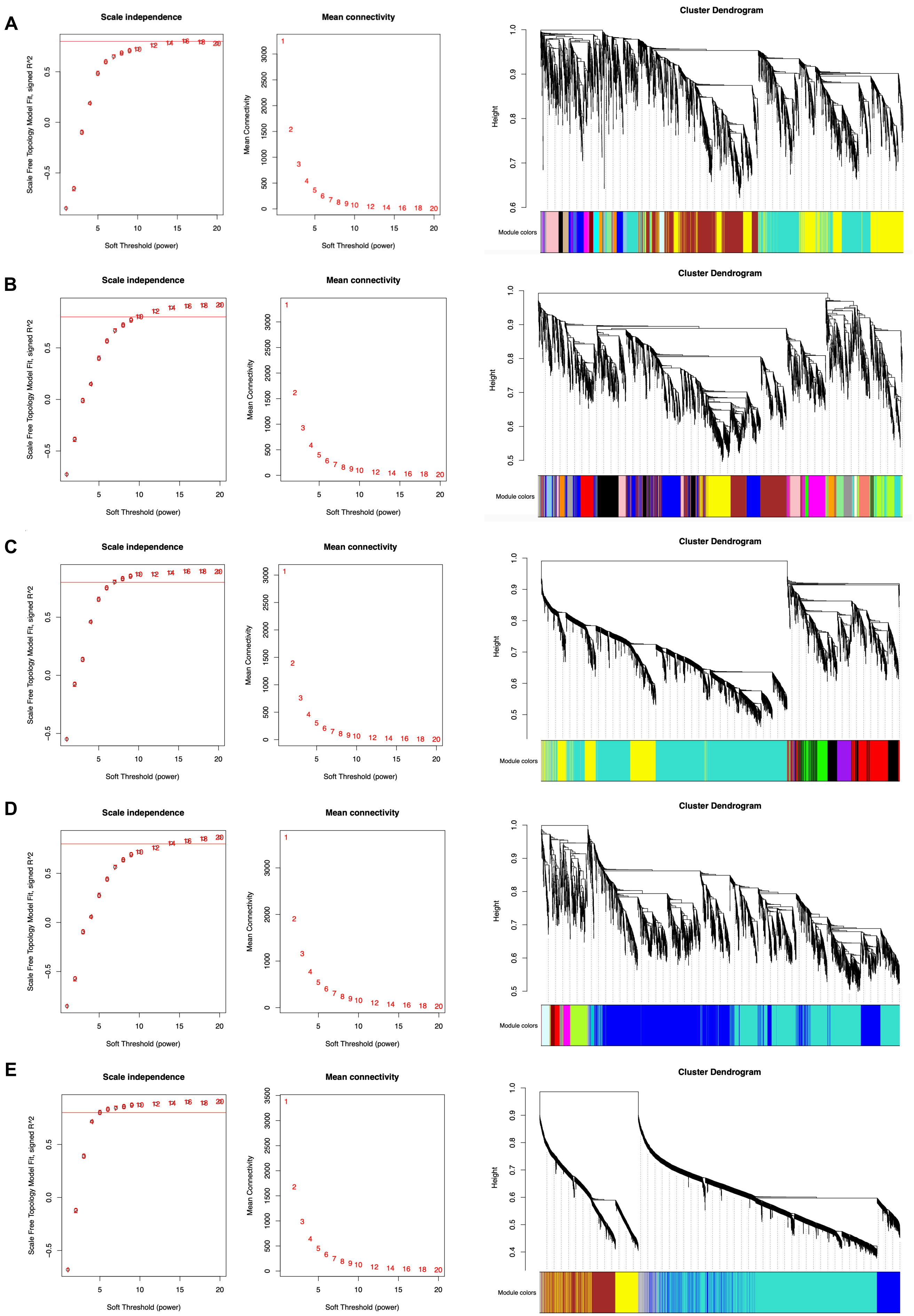
A
Scale independence
Scale Free Topology Model Fit, signed R^2
Soft Threshold (power)
Mean connectivity
Mean Connectivity
Soft Threshold (power)
Cluster Dendrogram
Height
Module colors
B
Scale independence
Scale Free Topology Model Fit, signed R^2
Soft Threshold (power)
Mean connectivity
Mean Connectivity
Soft Threshold (power)
Cluster Dendrogram
Height
Module colors
C
Scale independence
Scale Free Topology Model Fit, signed R^2
Soft Threshold (power)
Mean connectivity
Mean Connectivity
Soft Threshold (power)
Cluster Dendrogram
Height
Module colors
D
Scale independence
Scale Free Topology Model Fit, signed R^2
Soft Threshold (power)
Mean connectivity
Mean Connectivity
Soft Threshold (power)
Cluster Dendrogram
Height
Module colors
E
Scale independence
Scale Free Topology Model Fit, signed R^2
Soft Threshold (power)
Mean connectivity
Mean Connectivity
Soft Threshold (power)
Cluster Dendrogram
Height
Module colors

F

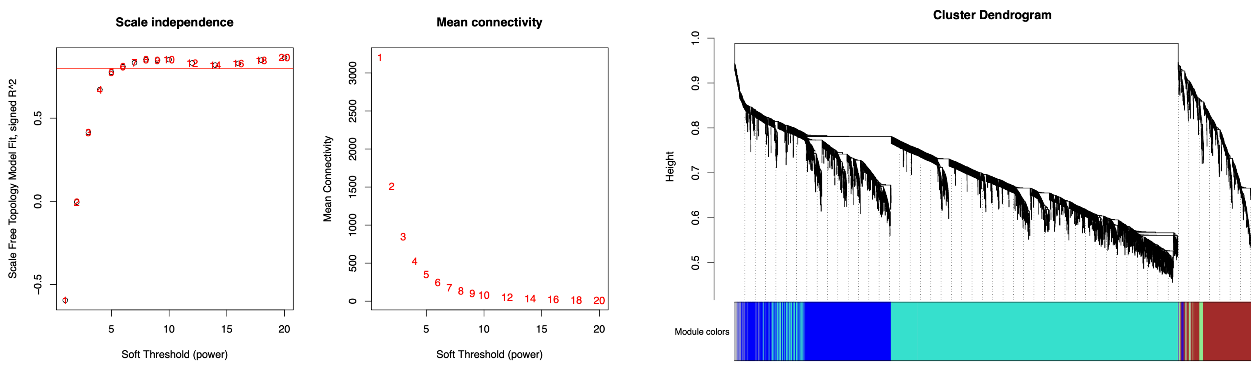


**Supplementary Figure 6. Determination of soft-threshold power and module detection in co-expression networks.**

Panels A (cattle), B (human), C (mouse), D (pig), E (rat), and F (marmoset) display analyses for each species. The left plots show the selection of soft-thresholding power to construct a scale-free topology, with scale independence (left) and mean connectivity (middle) plotted. The red line represents the threshold for scale-free topology ($R^2 > 0.8$), guiding the choice of power for network construction. The right plots in each panel show hierarchical clustering dendrograms of genes and their corresponding module assignments. Colored bars below the dendrograms indicate the identified modules.

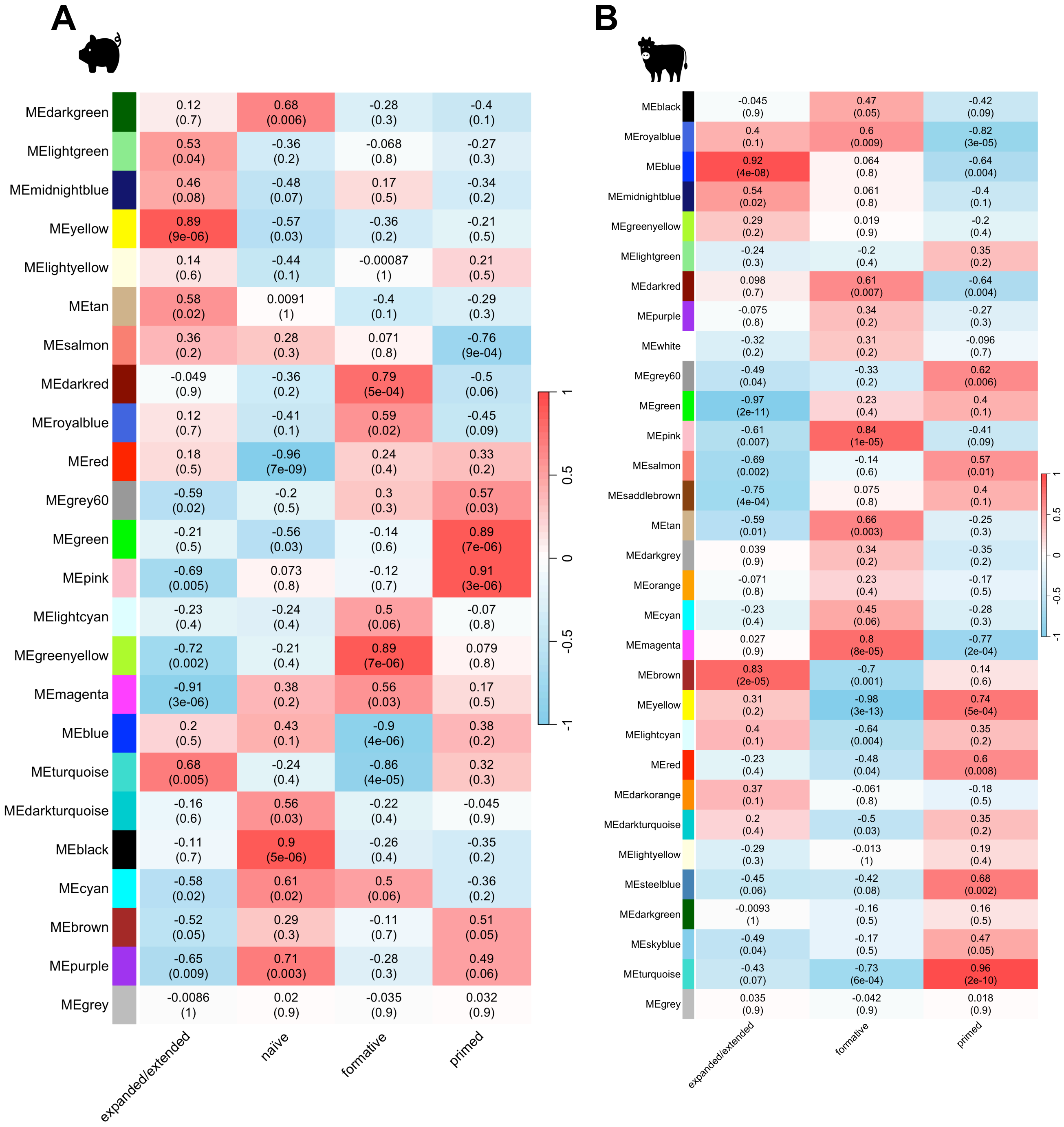
A
MEdarkgreen 0.12 (0.7) 0.68 (0.006) -0.28 (0.3) -0.4 (0.1)
MElightgreen 0.53 (0.04) -0.36 (0.2) -0.068 (0.8) -0.27 (0.3)
MEmidnightblue 0.46 (0.08) -0.48 (0.07) 0.17 (0.5) -0.34 (0.2)
MEyellow 0.89 (9e-06) -0.57 (0.03) -0.36 (0.2) -0.21 (0.5)
MElightyellow 0.14 (0.6) -0.44 (0.1) -0.00087 (1) 0.21 (0.5)
MEtan 0.58 (0.02) 0.0091 (1) -0.4 (0.1) -0.29 (0.3)
MEsalmon 0.36 (0.2) 0.28 (0.3) 0.071 (0.8) -0.76 (9e-04)
MEdarkred -0.049 (0.9) -0.36 (0.2) 0.79 (5e-04) -0.5 (0.06)
MEroyalblue 0.12 (0.7) -0.41 (0.1) 0.59 (0.02) -0.45 (0.09)
MEred 0.18 (0.5) -0.96 (7e-09) 0.24 (0.4) 0.33 (0.2)
MEgrey60 -0.59 (0.02) -0.2 (0.5) 0.3 (0.3) 0.57 (0.03)
MEgreen -0.21 (0.5) -0.56 (0.03) -0.14 (0.6) 0.89 (7e-06)
MEpink -0.69 (0.005) 0.073 (0.8) -0.12 (0.7) 0.91 (3e-06)
MElightcyan -0.23 (0.4) -0.24 (0.4) 0.5 (0.06) -0.07 (0.8)
MEgreenyellow -0.72 (0.002) -0.21 (0.4) 0.89 (7e-06) 0.079 (0.8)
MEmagenta -0.91 (3e-06) 0.38 (0.2) 0.56 (0.03) 0.17 (0.5)
MEblue 0.2 (0.5) 0.43 (0.1) -0.9 (4e-06) 0.38 (0.2)
MEturquoise 0.68 (0.005) -0.24 (0.4) -0.86 (4e-05) 0.32 (0.3)
MEdarkturquoise -0.16 (0.6) 0.56 (0.03) -0.22 (0.4) -0.045 (0.9)
MEblack -0.11 (0.7) 0.9 (5e-06) -0.26 (0.4) -0.35 (0.2)
MEcyan -0.58 (0.02) 0.61 (0.02) 0.5 (0.06) -0.36 (0.2)
MEbrown -0.52 (0.05) 0.29 (0.3) -0.11 (0.7) 0.51 (0.05)
MEpurple -0.65 (0.009) 0.71 (0.003) -0.28 (0.3) 0.49 (0.06)
MEgrey -0.0086 (1) 0.02 (0.9) -0.035 (0.9) 0.032 (0.9)
expanded/extended
naïve
formative
primed
1
0.5
0
-0.5
-1
B
MEblack -0.045 (0.9) 0.47 (0.05) -0.42 (0.09)
MEroyalblue 0.4 (0.1) 0.6 (0.009) -0.82 (3e-05)
MEblue 0.92 (4e-08) 0.064 (0.8) -0.64 (0.004)
MEmidnightblue 0.54 (0.02) 0.061 (0.8) -0.4 (0.1)
MEgreenyellow 0.29 (0.2) 0.019 (0.9) -0.2 (0.4)
MElightgreen -0.24 (0.3) -0.2 (0.4) 0.35 (0.2)
MEdarkred 0.098 (0.7) 0.61 (0.007) -0.64 (0.004)
MEpurple -0.075 (0.8) 0.34 (0.2) -0.27 (0.3)
MEwhite -0.32 (0.2) 0.31 (0.2) -0.096 (0.7)
MEgrey60 -0.49 (0.04) -0.33 (0.2) 0.62 (0.006)
MEgreen -0.97 (2e-11) 0.23 (0.4) 0.4 (0.1)
MEpink -0.61 (0.007) 0.84 (1e-05) -0.41 (0.09)
MEsalmon -0.69 (0.002) -0.14 (0.6) 0.57 (0.01)
MEsaddlebrown -0.75 (4e-04) 0.075 (0.8) 0.4 (0.1)
MEtan -0.59 (0.01) 0.66 (0.003) -0.25 (0.3)
MEdarkgrey 0.039 (0.9) 0.34 (0.2) -0.35 (0.2)
MEorange -0.071 (0.8) 0.23 (0.4) -0.17 (0.5)
MEcyan -0.23 (0.4) 0.45 (0.06) -0.28 (0.3)
MEmagenta 0.027 (0.9) 0.8 (8e-05) -0.77 (2e-04)
MEbrown 0.83 (2e-05) -0.7 (0.001) 0.14 (0.6)
MEyellow 0.31 (0.2) -0.98 (3e-13) 0.74 (5e-04)
MElightcyan 0.4 (0.1) -0.64 (0.004) 0.35 (0.2)
MEred -0.23 (0.4) -0.48 (0.04) 0.6 (0.008)
MEdarkorange 0.37 (0.1) -0.061 (0.8) -0.18 (0.5)
MEdarkturquoise 0.2 (0.4) -0.5 (0.03) 0.35 (0.2)
MElightyellow -0.29 (0.3) -0.013 (1) 0.19 (0.4)
MEsteelblue -0.45 (0.06) -0.42 (0.08) 0.68 (0.002)
MEdarkgreen -0.0093 (1) -0.16 (0.5) 0.16 (0.5)
MEskyblue -0.49 (0.04) -0.17 (0.5) 0.47 (0.05)
MEturquoise -0.43 (0.07) -0.73 (6e-04) 0.96 (2e-10)
MEgrey 0.035 (0.9) -0.042 (0.9) 0.018 (0.9)
expanded/extended
formative
primed
1
0.5
0
-0.5
-1

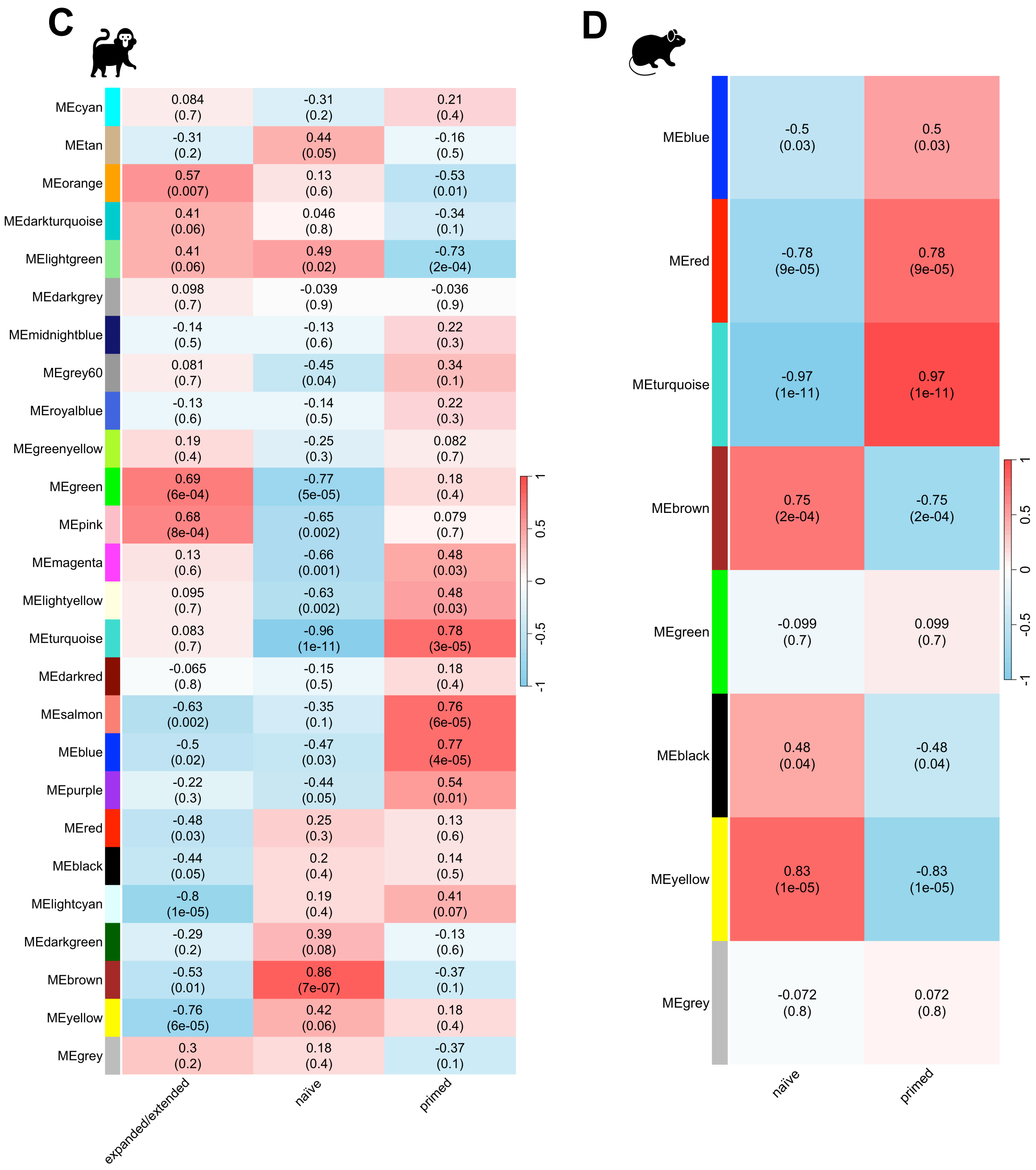


**Supplementary Figure 7. Module-trait associations in co-expression networks across species.**

WGCNA analysis was performed separately for each species using the soft threshold determined in Figure S6. Results are shown for pig (A), cattle (B), marmoset (C), and rat (D).

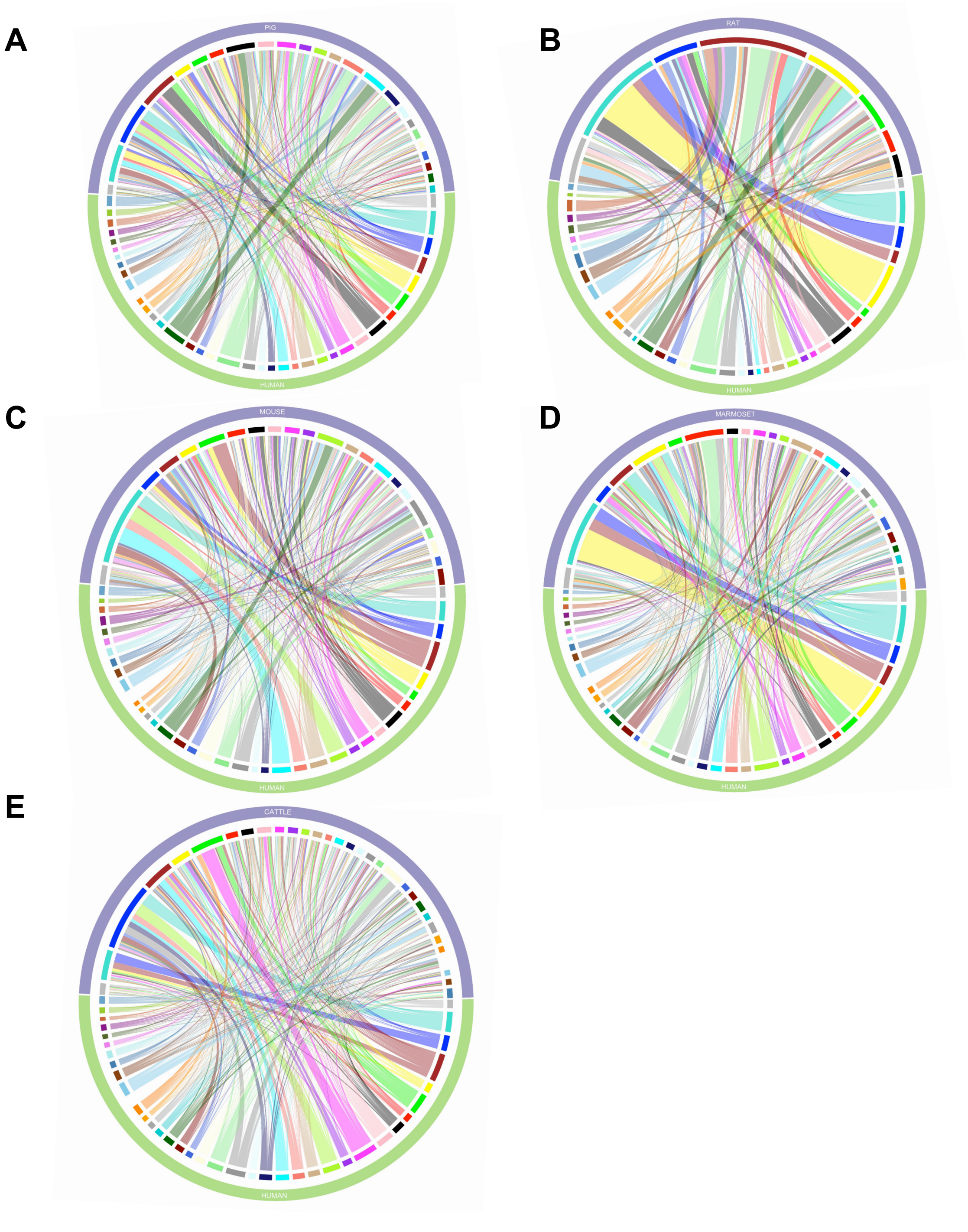


**Supplementary Figure 8. Preservation of modules between the networks of other species and the human network.**

This figure illustrates the preservation of human network modules in the networks of other species. The thickness of the connecting bands represents the –log10(P) value, indicating the strength of preservation. The bar colors on the inner circle correspond to the module names. Panels show results for pig (A), rat (B), mouse (C), marmoset (D), and cattle (E).

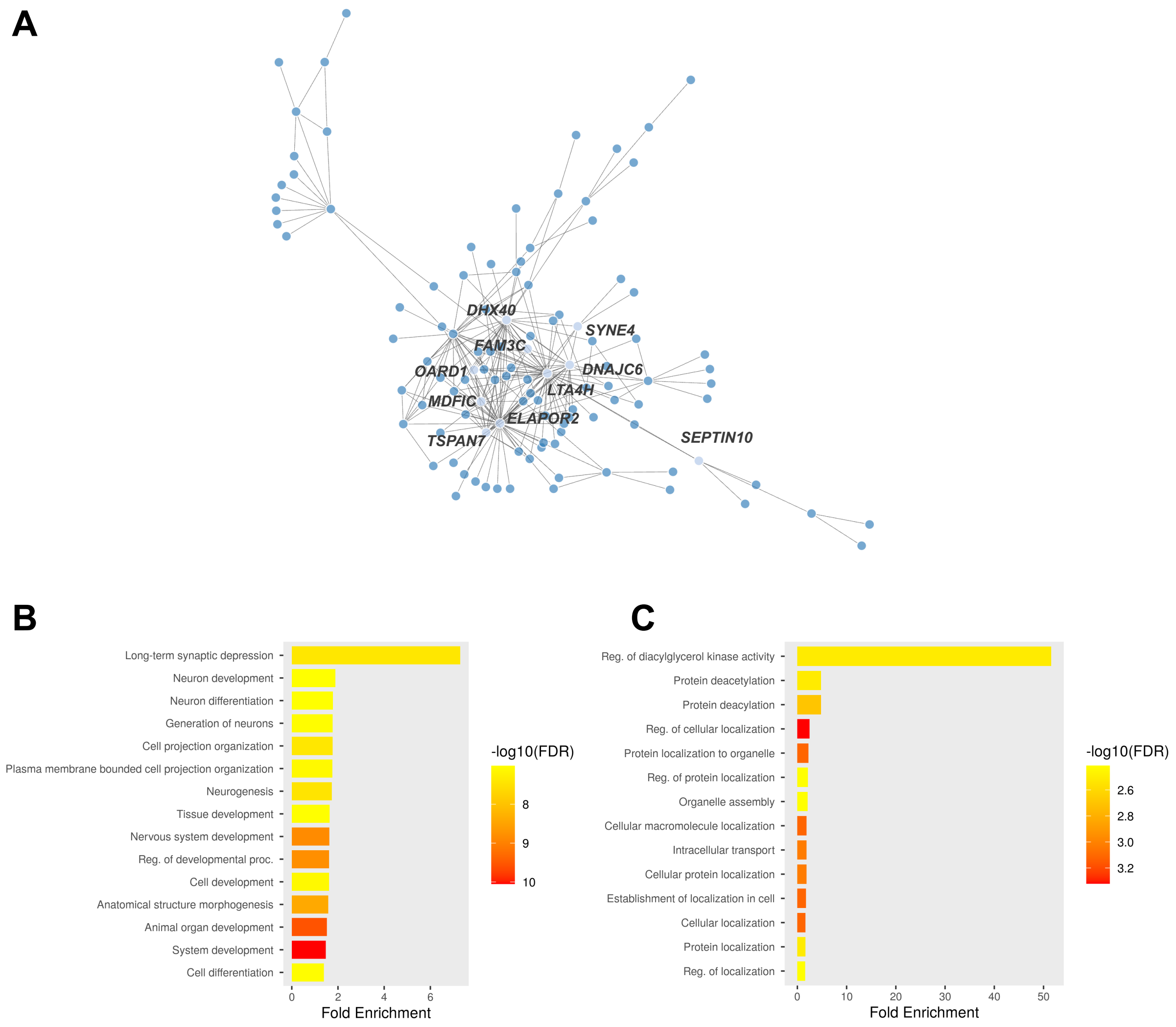


**Supplementary Figure 9. Overview of co-expression networks and functional enrichment analyses for human black and blue modules.**

A. Gene co-expression network of the human black module constructed using WGCNA, with top hub genes highlighted.

B. GO:BP analysis showing pathways enriched by genes in the human blue module.

C. GO:BP analysis showing pathways enriched by genes in the human black module.

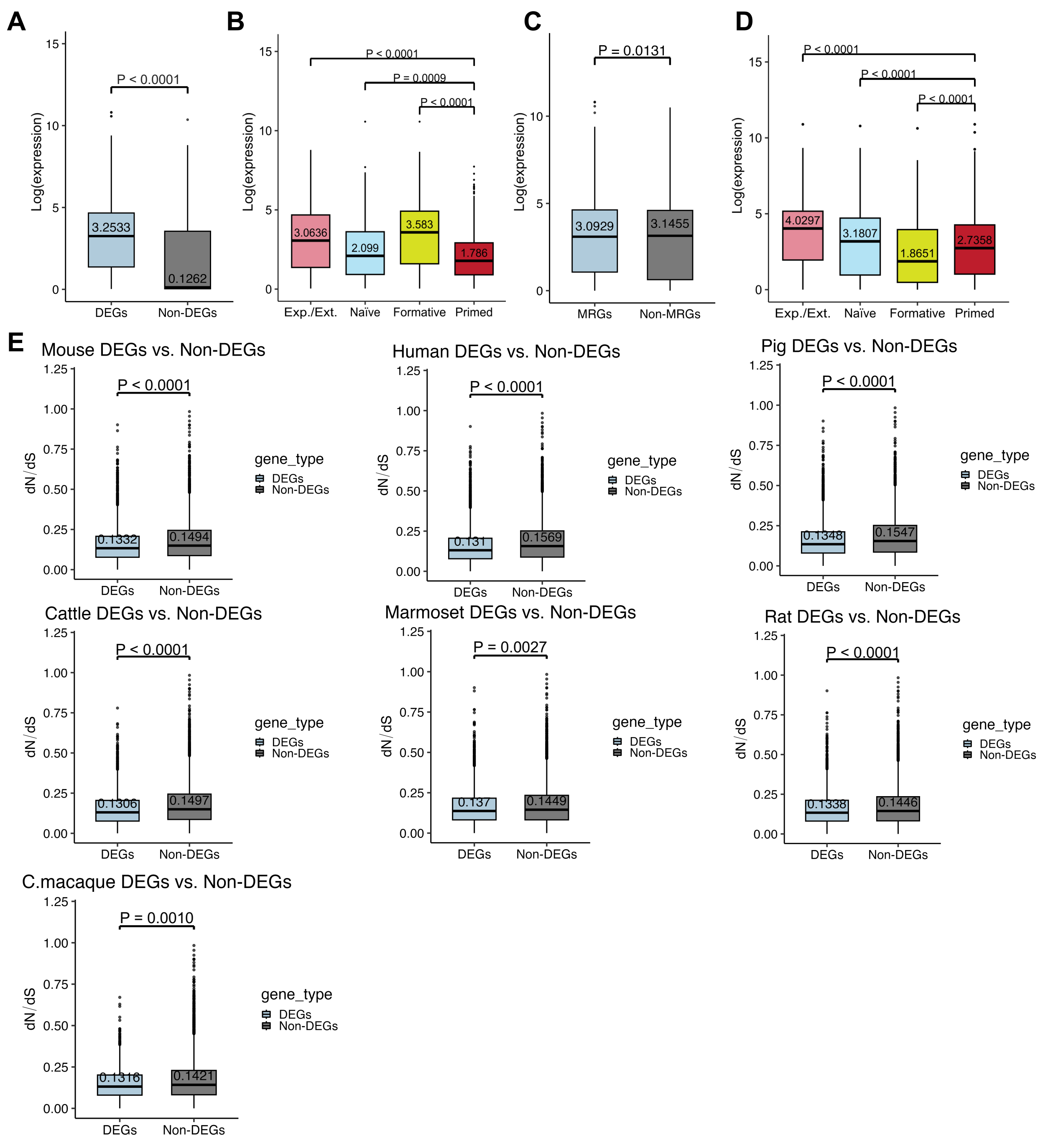

A
B
C
D
E
Log(expression)
P < 0.0001
3.2533
0.1262
DEGs
Non-DEGs
P < 0.0001
P = 0.0009
P < 0.0001
3.0636
2.099
3.583
1.786
Exp./Ext.
Naïve
Formative
Primed
P = 0.0131
3.0929
3.1455
MRGs
Non-MRGs
4.0297
3.1807
1.8651
2.7358
Mouse DEGs vs. Non-DEGs
0.1332
0.1494
Human DEGs vs. Non-DEGs
0.131
0.1569
Pig DEGs vs. Non-DEGs
0.1348
0.1547
Cattle DEGs vs. Non-DEGs
0.1306
0.1497
Marmoset DEGs vs. Non-DEGs
P = 0.0027
0.137
0.1449
Rat DEGs vs. Non-DEGs
0.1338
0.1446
C.macaque DEGs vs. Non-DEGs
P = 0.0010
0.1316
0.1421
dN/dS
gene_type
DEGs
Non-DEGs

F

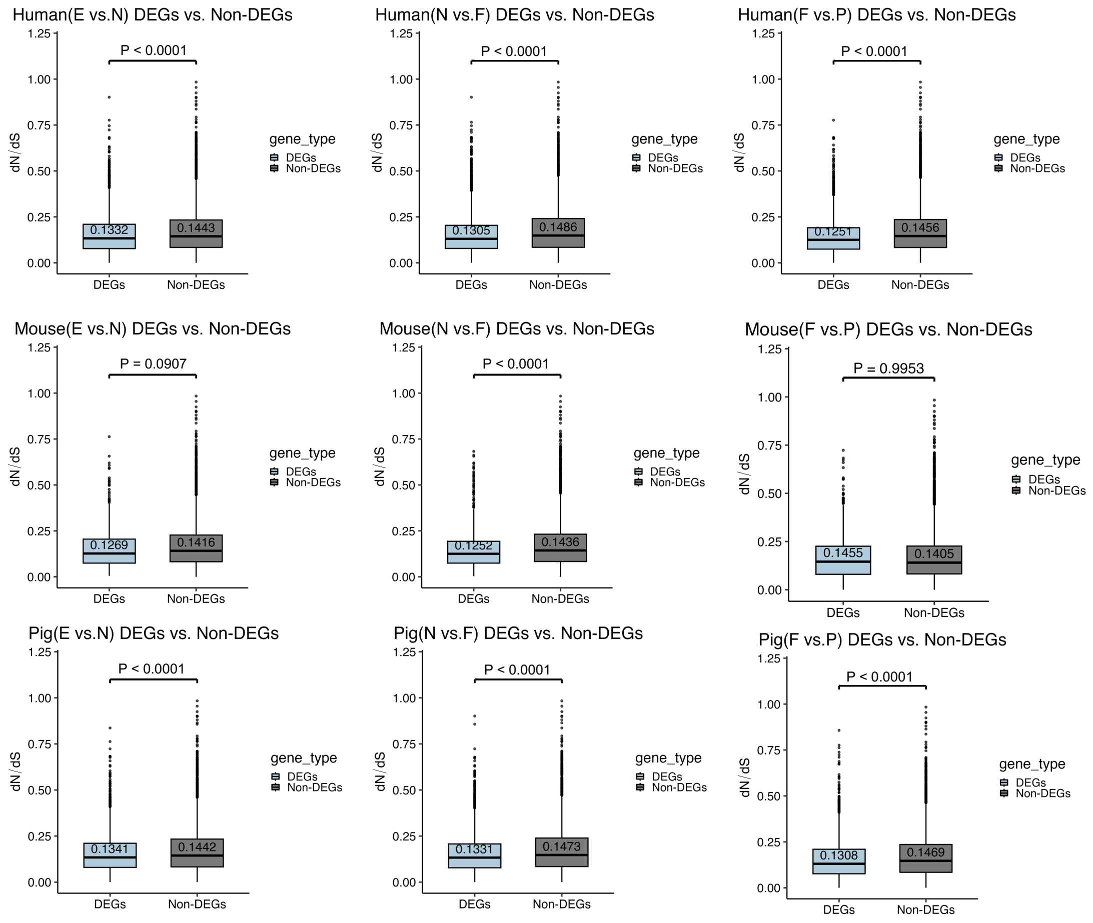


**Supplementary Figure 10. Comparative Analysis of Gene Expression and Evolutionary Constraints Across Pluripotency States.**

**A–D. Box plots showing mean expression values for various gene sets:** DEGs versus non-DEGs (A), state-specific genes identified through DEG analysis (B), module-associated genes (MRGs) identified by WGCNA as related to any pluripotency state versus non-MRGs (C), and genes in WGCNA modules associated with specific states (D).

**E–F. Box plots showing dN/dS ratios for DEGs versus non-DEGs:** all DEGs versus non-DEGs across nascent pluripotency states for each species (E) and DEGs versus non-DEGs from separate DEG comparisons of nascent pluripotency states for individual species (F).
